# Hydration free energy is a significant predictor of globular protein incorporation into condensates

**DOI:** 10.1101/2025.07.03.663089

**Authors:** Sophie Anderson, Malcolm Harrison, Gregory L Dignon

**Author notes:** These authors contributed equally to this work (Sophie Anderson, Malcolm Harrison).

## Abstract

Membraneless organelles (MLOs) are assemblies of biomolecules, which function without a dividing lipid membrane in a cellular environment. These MLOs, termed biomolecular condensates, are commonly formed by the thermodynamic process of liquid-liquid phase separation (LLPS) and assembly of large numbers of proteins, nucleic acids and co-solvent molecules. Within MLOs, certain biomolecule types are particularly causative of phase separation, and are termed “scaffolds” as they provide the major driving forces for self-assembly. Other molecules that are present in a condensate, but are less causative than the scaffold molecules are termed “clients”. Much effort has recently decoded many of the molecular interactions underlying LLPS in search of predicting equilibrium concentrations and materials properties of condensates. In this work, we provide a simple computational approach to predict the partitioning of globular protein clients into condensates primarily composed of disordered protein scaffolds. Specifically, we use multiple methods to calculate hydration free energy of a series of globular proteins, and find that hydration free energy is relatively well-correlated with the partition coefficient of these proteins into condensates. We then provide a comparison of different hydration free energy predictors and discuss why some may provide a more accurate prediction of partitioning. Finally, we discuss the shortcomings of hydration free energy as a predictor by identifying other possible confounding factors such as specific interactions, charge matching, and differential solvation inside a condensate, which will aid in making more robust predictions in future studies trained on more diverse data sets.

**Significance Statement:** Understanding the extent to which molecules can partition into biomolecular condensates is crucial for deciphering cellular organization and function. This study introduces a simple computational approach to predict the partitioning of globular proteins into condensates using hydration free energy as a key predictor, which can be calculated from static globular protein structures. By comparing different hydration free energy predictors, we also higlight the limitations of relying solely on this hydration free energy, emphasizing the need to consider other molecular factors. We finally analyze the shortcomings of the hydration free energy precitions, and discuss other factors that contribute to partitioning of clients into condensates, namely specific interactions and net charge of the scaffold molecules, and different properties of water in the condensate.

## Introduction

Biomolecular condensates, or membraneless organelles (MLOs), are cellular compartments made up of self-assembled biomolecules that lack a dividing lipid membrane as is present in membrane-bound organelles[1, 2]. Many biomolecular condensates are formed through the process of Liquid-Liquid Phase Separation (LLPS), in which a homogeneous solution of macromolecules separates into two or more liquid-like phases having distinct molecular compositions[3, 4, 5]. The self-association of molecules involved in driving LLPS is predominantly due to weak multivalent interactions between proteins and nucleic acids, associating with each other to form a “dense” phase and isolating from the surrounding “dilute” phase[6, 7, 8]. The interior of a condensate differs from that of its surroundings through various emergent properties including surface tension, viscoelasticity, and differential solvation of small molecules[9, 10]. Biomolecular condensates coordinate and execute specific functions integral to various biological processes, including DNA repair, cellular stress response, filtration, and signal transduction[11, 12, 13, 14]. Dysregulated biomolecular condensates are associated with the development and progression of cancer, as well as neurological diseases such as ALS[15, 16, 17, 18].

Most MLOs contain dozens to hundreds of types of molecules, including proteins, nucleic acids, small molecules and solvent[19, 20, 21, 22, 23]. Molecules that are particularly causative of phase separation through attractive interactions are termed scaffolds, while other components that are less causative, but still incorporate into condensates are termed clients[21, 4, 24]. The identity and materials properties of a condensate are largely linked to the composition and concentration of molecular components and their interconnectedness. Therefore it is essential to have a good understanding of what driving forces control composition of condensates for characterization of condensate function and materials properties, and for design of new clients, and condensate-interacting molecules[25, 4, 24, 22].

Many studies have helped decode molecular grammar of phase separation, particularly focusing on sequence and composition of intrinsically disordered regions (IDRs) which are common in phase-separating proteins[26, 27, 28, 29, 30, 31]. However, many condensates include partially folded, or completely folded globular proteins, which cannot be fully described based on their amino acid sequence[32, 33]. It is useful to establish simple and generalizable metrics to quantify and predict the partitioning of globular proteins into condensates, as this enables a better understanding of the molecular determinants that govern condensate composition and function, and facilitates the development of predictive models for biological and biomedical applications.

Multicomponent phase separation can occur through several distinct modes, including cooperative cophase separation, scaffold-client cophase separation, cross-interaction-driven cophase separation, and exclusive phase separation[4, 25, 34, 35]. Among these, scaffold-client systems are particularly widespread and experimentally tractable, making them valuable models for investigating the factors that influence client protein incorporation[19, 36, 22]. However, the highly dynamic and disordered nature of condensate interiors presents challenges in identifying which specific protein interactions are most important for phase separation and client partitioning. Therefore, development of predictive metrics can aid in elucidating the underlying physical principles that determine condensate composition and inform future studies of protein behavior in these environments.

In this work, we derive a thermodynamic cycle that relates hydration free energy to the transfer free energy of GFP variants partitioning into FG nucleoporin condensates, providing a quantitative model for scaffold-client systems. We assess the correlation between the experimentally measured partition coefficients (P) of green fluorescent protein (GFP) variants and their computed hydration free energies (Δ*F_hyd_*), finding that hydration free energy is a strong predictor of partitioning behavior. We then systematically evaluate several computational models for estimating hydration free energy, finding that static structure-based approaches such as the hydrophibic intensity patch (hi-patch) method and effective energy function 1 (EEF1) are decently predictive of experimental partition coefficients while being computationally inexpensive, and relatively easy to use. Finally, we conduct an analysis of possible sources of deviation from these predictions, highlighting additional factors such as the scaffold molecule’s physicochemical properties, and specific scaffold-client interactions, each of which have an influence on partitioning.

## Results & Discussion

### Thermodynamic cycle of solvation and hydration

Hydration free energy (Δ*F_hyd_*) is a measure of the change in free energy of a molecule from vacuum to pure water solvent (Fig. 1), and will generally be negative for soluble molecules[37]. Hydration free energy is commonly approximated by the transfer from oil to water, since oil represents a hydrophobic environment[38]. Similarly, the transfer free energy of a molecule partitioning into a condensate can be mathematically defined as the difference between the free energy of a molecule inside and outside a condensate, and is logarithmically related to the molecule’s parition coefficient (P):

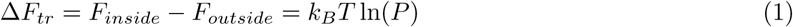

where *P* = *c_cond_/c_sat_*is the ratio of a molecule’s concentration inside the condensate (c*_cond_*) to it’s concentration outside the condensate (c*_sat_*). For scaffolds, this tends to be a large number on the order of tens to thousands, while for clients it can be more moderate, or even less than 1 for a molecule that is excluded from a condensate. Notably, in multicomponent systems, the partition coefficient and saturation concentration is dependent on the total overall concentration of each component in the system[39, 4, 24]. Various experimental methods are capable of measuring partition coefficients, and thus can be used to calculate transfer free energy of a client protein into a condensate composed of a separate scaffold protein[23, 20]. In experiments where the condensate is primarily composed of scaffold, and the external phase has a very low scaffold concentration, there should be minimal effect on the client’s partition coefficient, as it is similar to the scenario described by the definition of Δ*F_tr_*.

**Figure 1:**
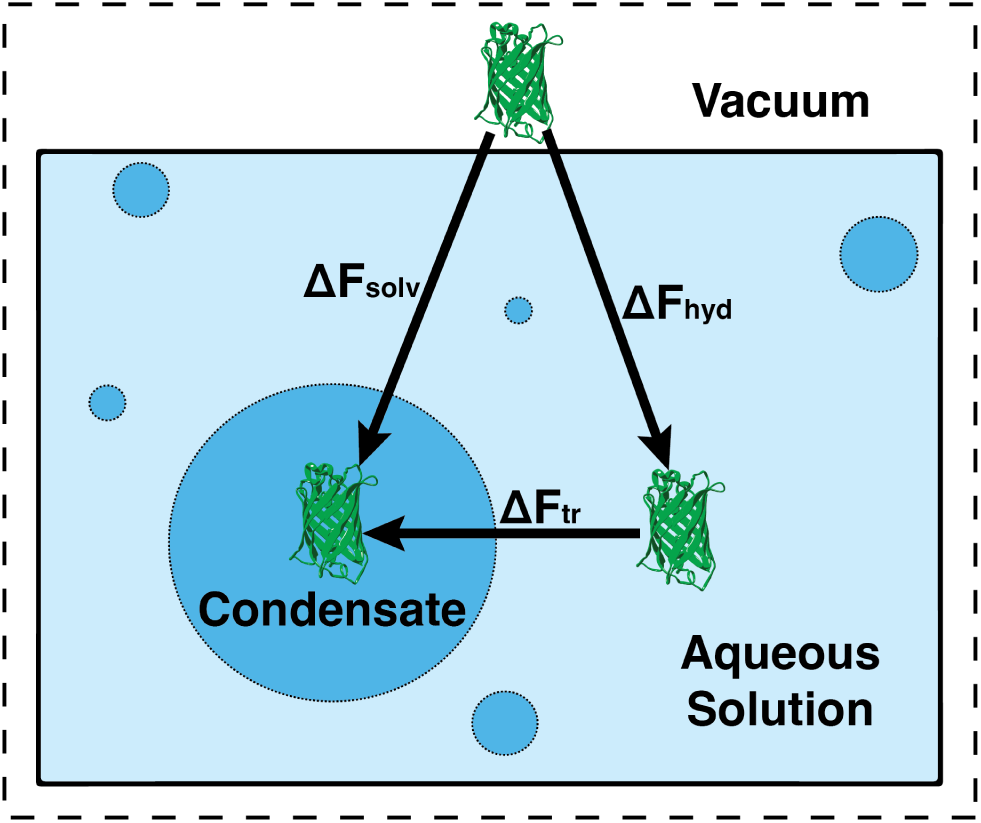
A simple thermodynamic cycle to describe transfer free energy of protein between vacuum, low concentration aqueous solution, and condensate environments.

Notably, this ΔF*_tr_* represents the free energy required to transfer between two aqueous phases, specifically between aqueous surroundings and a water-containing protein-rich condensate, and could be expected to differ significantly from oil-water transfer free energy of the same molecule. However, recent work has shown that oil-water transfer is a good predictor of small molecule partitioning into condensates, perhaps due to the differential solvent behavior of water inside a condensate[10]. To fully describe the transfer free energy of a protein into a condensate and relate it to hydration free energy, we devise a thermodynamic cycle, distinguishing between transfer from vacuum to pure water, vacuum to condensate, and the difference between these two being equivalent to the transfer free energy (Fig. 1).

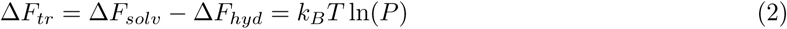

We note that the value of Δ*F_solv_* is more challenging to obtain directly, but can be calculated indirectly through the thermodynamic cycle if Δ*F_tr_* and Δ*F_hyd_* are determined indivicually. This value would indicate the free energy change upon insertion to a condensate from vacuum, which is of little practical value. Thus we focus primarily on the relation between Δ*F_hyd_* and Δ*F_tr_* for the remainder of this paper, and only invoke Δ*F_solv_* to explain deviations in the data. Notably, values of Δ*F_tr_* are of much smaller magnitude than Δ*F_hyd_*or Δ*F_solv_*due to the similarity of water to condenesate interior as compared to vacuum.

### Relation between hydration free energy and partition coefficients of GFP variants

As a model system, we use data from Frey et al. who developed a series of green fluorescent protein (GFP) variants with the same folded structure, but different surface properties. Namely, they mutated eight sites on the surface of the protein, all to one particular amino acid, for 17 of the 20 canonical amino acids (Table S1). Using two different FG nucleoporin proteins (FG Nups; Nup116 and MacNup98A) as scaffolds, they quantified the partitioning of the 17 GFP variants into condensates of each scaffold, resulting in 34 total measurements of partition coefficients[20]. We can convert these measurements of partition coefficients to transfer free energies using Eq. 2. We find that the transfer free energy of the globular proteins into condensates can vary from most incorporated -14.5 kJ/mol (8W) to most excluded, 7.0 kJ/mol (8K) at 298K (Table S2). We note that the values of Δ*F_hyd_* derived from experiments correlate well with the hydropathy values reported by Urry et al., and used in coarse-grained simulations of condensates (Fig. S1)[40, 41].

To approximate the Δ*F_tr_* of each GFP variant into the Nup condensates, we calculated the Δ*F_hyd_* using static structures of each globular GFP variant using hi-patch. Hi-patch is a method designed to identify hydrophobic patches on protein surfaces, based on parameters from the EEF1 implicit solvation model, and correcting each atom’s hydration based on its neighboring atoms[42, 43]. This approach is motivated by observations that hydropathy is influenced not only by presence of nonpolar atoms, but also by patchiness of nonpolar atoms, having many within close proximity to each other[44, 45]. The hi-patch method calculates multiple surface geometry-based features including number of hydrophobic patches, the surface area of each patch, and the average hydropathy of each patch, among others (Fig. S2, Table S3). We find that of these properties, the absolute hydration free energy, ΔF*_hyd_* is the best predictor of ΔF*_tr_* (Fig. 2, S3). We find that more negative values of Δ*F_hyd_* correspond to greater partitioning of the client into the condensate, highlighting an inverse correlation between Δ*F_hyd_* and Δ*F_tr_*. This is reasonable because the condensate environment, while still aqueous, has a smaller water content than the exterior, making more soluble proteins prefer the aqueous exterior to the condensate interior. Notably, the hydration free energy of only the hydrophobic patches on the protein surface is also a reasonable predictor for the partition coefficients, as well as the total number of patches, and the aromaticity of the surface (Fig. S3). Of these, the aromaticity metric is the least redundant with total hydration free energy, having a covariance of 0.6 with total Δ*F_hyd_* (Fig. S2). Such additional features may be added to the predictive model, but risk overfitting the model in absence of a larger training data set.

**Figure 2:**
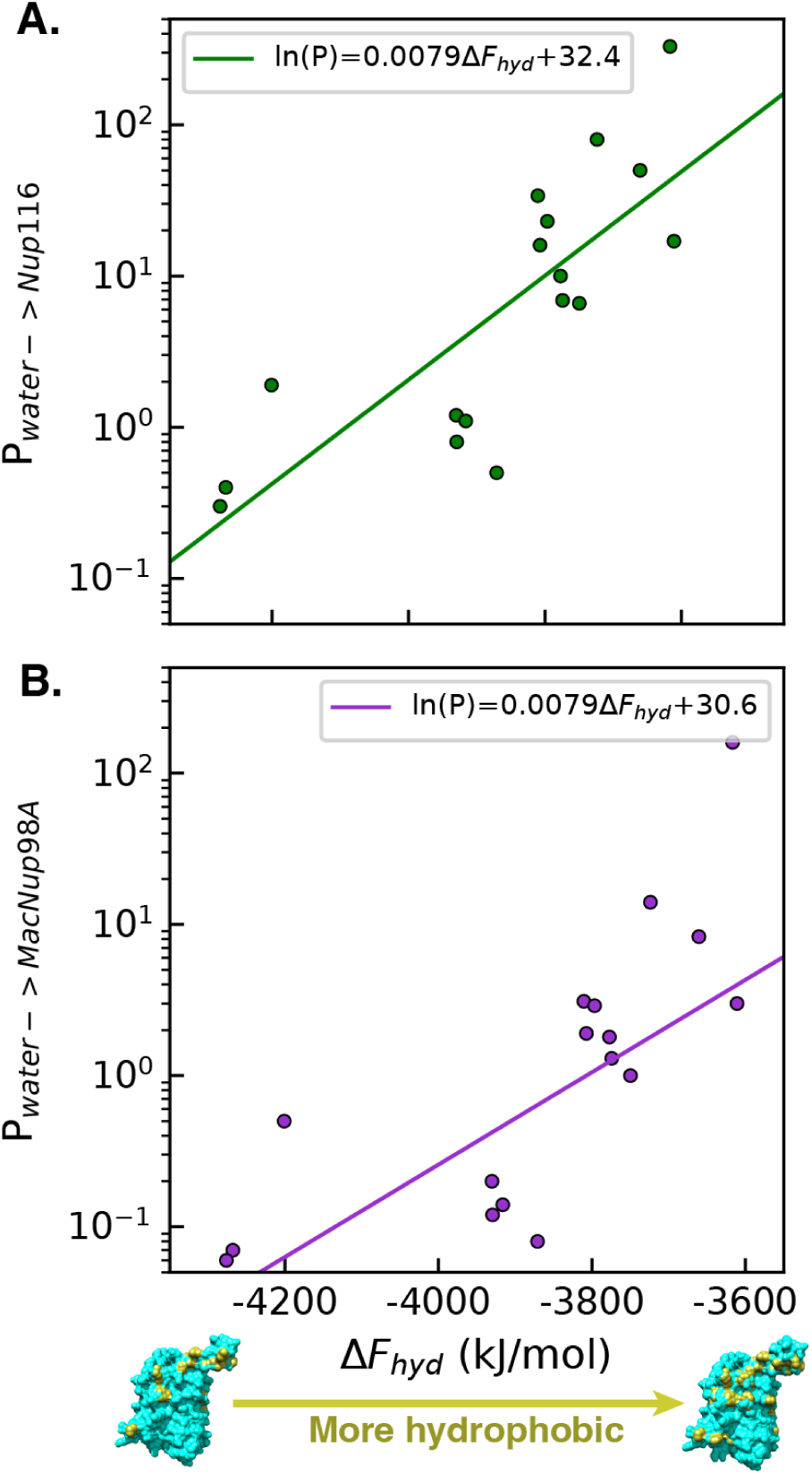
Correlation of computationally-calculated Δ*F_hyd_* and experimentally-measured partition coefficient for 17 variants of GFP into A) Nup116, and B) MacNup98A. Renderings show least hydrophobic and most hydrophobic GFP variants visualized using hi-patch per-atom hydropathy output.

Thus, we conclude that predicted Δ*F_hyd_* of a protein is a decent predictor of partitioning of a globular protein into a condensate, but does not account for all factors. Another consideration is that the relationship between Δ*F_hyd_* and ln(P) is quantitatively different between Nup116 and MacNup98A. While the Δ*F_hyd_*of each globular protein is independent of the scaffold molecule, the partition coefficients will usually be different, thus resulting in different quantitative fits. Since Nup116 is longer than MacNup98A it has greater valency, and number of aromatic residues (Table S4), and thus it stands to reason that the clients would be more prone to partition into their condensates due to greater interaction propensities with the scaffolds[46]. Within this data set, the two scaffold molecules have relatively similar composition (Table S4-5), and so there is not a very large difference in partitioning of clients into these condensates. It is plausible that partitioning of these same clients into condensates of compositionally distinct scaffold proteins would result in more significant deviation from this behavior[47, 22].

### Comparison of hydration models for predicting partitioning

While the hi-patch method provides decent predictive capabilities of GFP partitioning into Nup condensates, we ask the question whether other hydration models are more appropriate for this problem. Thus, we carried out ΔF*_hyd_* calculations using two additional static methods, applied directly to static structures, as well as three dynamic approaches that carry out molecular dynamics (MD) simulations of the protein to capture fluctuations of the structure and surface of the protein.

Table 1 shows the different hydration models used and the correlation to the log of the experimentally-measured partition coefficients. The models chosen for these ΔF*_hyd_* calculations all vary in how they approximate solvation free energy and what they include or neglect to balance accuracy with computational power. CHARMM EEF1 is a great base for calculating ΔF*_hyd_*, and in the 25 years since its publication several advancements in computation speed have allowed for more accurate approximations, such as fast surface area calculations and the ability to use environmental corrections and account for neighboring atoms affecting solubility, as in the updates applied in the hi-patch method[42, 48]. The APBS model does not explicitly account for ion-solvent interactions and models the solvent as a dielectric continuum, which can affect the accuracy of the approximate hydration free energy[49]. The GBNeck model approximates the Generalized Born implicit solvent model to be able to obtain a solution faster for use with MD simulations, which may result in errors from the true value of the change in ΔF*_hyd_*[50].

**Table 1:** Quality of agreement between different hydration models and experimental partition coefficients. The hi-patch algorithm, CHARMM EEF1 and ABPS were applied to static structures. Note that GBNeck and hi-patch traj implicit methods are applied to the same trajectory, just using different hydration models.

| Method | Input | Nup116 $R^2$ | MacNup98A $R^2$ |
| --- | --- | --- | --- |
| hi-patch | static | 0.6334 | 0.5752 |
| CHARMM EEF1 | static | 0.3981 | 0.3166 |
| ABPS | static | 0.1289 | 0.1185 |
| GBNeck | dynamic | 0.1015 | 0.0919 |
| hi-patch traj explicit | dynamic | 0.4759 | 0.4154 |
| hi-patch traj implicit | dynamic | 0.3815 | 0.2721 |

To assess the effect that dynamic structure change has on ΔF*_hyd_*, molecular dynamics (MD) simulations were performed using both the GBNeck implicit solvent model, as well as explicit solvent simulations in TIP3P water[51]. We directly calculated the hydration free energy of the GFP molecules from GBNeck as reported by the implicit solvent model and calculated the average over the trajectory. This resulted in the poorest prediction of experimentally measured partition coefficients. We thus applied the hi-patch algorithm to the trajectory generated by both the GBNeck implicit solvent model, and the explicit solvent simulation, taking the average of all frames, and finding that it provided better predictions for experimental partition coefficients. This indicates that the APBS and GBNeck hydration models struggle to report qualitatively accurate values for hydration free energy between different similar proteins, while EEF1 and hi-patch provide the most useful qualitative predictions.

Interestingly, the hi-patch method provided the best predictions when applied to only a static structure, and weaker predictions when applied to a dynamic trajectory from MD simulations. The reason for this could be due to weaknesses in the conformational sampling of the MD simulations, or perhaps due to different populations of states occurring within the condensate that would not be accounted for in the bulk aqueous MD simulations. Out of all of the static and dynamic methods used to predict ΔF*_hyd_*, hi-patch had the strongest linear correlation between ΔF*_hyd_* and ln(P) (Fig. 2; Table 1). Furthermore, the hi-patch calculation can be carried out nearly instantaneously, and only requires a static 3D structure of the protein, making it accessible to implement without needing lots of computational setup and large amounts of computing power. It may be beneficial to attempt more rigorous quantitative calculations of hydration free energy[52, 37]. However, it is unclear whether the benefits of additional accuracy will be sufficient to account for significantly larger computational cost. Future work could evaluate additional methods, but would benefit from a larger and more diverse set of globular client proteins as a training set. From the current data set, we can still hypothesize possible causes for error in the predictions from hi-patch-calculated Δ*F_hyd_*.

### Specific interactions and differential solvation driving partitioning

We hypothesize three potential sources of error from our predictions as 1) specific interactions between the surface residues of GFP and the residues in the scaffold protein, 2) the matching of charges in the condensed phase, and 3) the altered solvent environment inside the condensate[47, 10, 53]. To explore this, we compute the error of predictions from the logarithmic fit between Δ*F_hyd_* and P from experimental results as Δ*F_tr_* Error = *RT* ln(*P_actual_ − P_predict_*) (Fig. 3).

**Figure 3:**
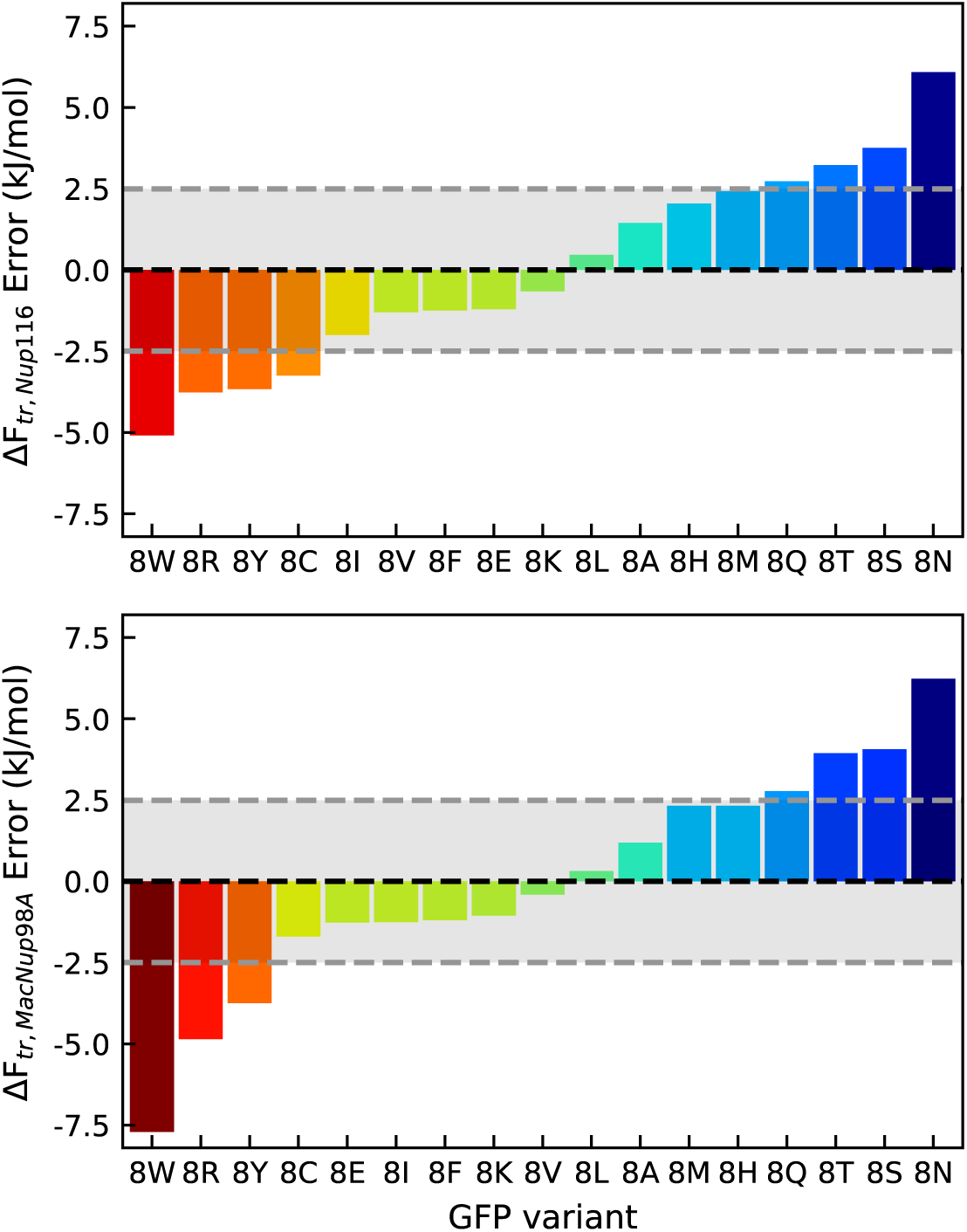
Sorted ΔF*_tr_* values comparing predicted partitioning, to experimental measurements for A) Nup116 and B) MacNup98A. Sorting was done to display order of most underpredicted to most overpredicted. Grey boxes are drawn around values of *±* 1 k*_B_*T.

Related to the question of specific interactions, we find that GFP variants decorated by tyrosine (Y), tryptophan (W), and arginine (R) residues are systematically underpredicted by our hydration-based model (Fig. 3). These residues, particularly R and Y have been identified as highly causative of phase separation in disordered proteins[26, 54] owing to their sticker-like behavior in the theory of associative polymers[55, 56]. This can be attributed to their unique interaction propensities since Y and W are aromatic and likely engage in *π*–*π* stacking with aromatic groups in FG nucleoporins, while R is known to participate in strong cation-*π* interactions with aromatic residues[57, 26, 54]. The guanidinium group of R is especially effective at such interactions, as it can simultaneously form hydrogen bonds and cation-*π* interactions, which may account for the greater underprediction of its partitioning relative to other positively charged residues[54]. We note that the amino acid composition, and particularly the aromatic fraction of both scaffold molecules is similar (Tables S4-5), so the extent of specific interactions confounding the purely hydration free energy-based prediction is likely to be similar between both cases. As such, we find both Nup116 and MacNup98A partition coefficients are underpredicted to similar degrees for each of these amino acids. We also find that W and Y are the most underpredicted clients for all hydration models tested in this work (Figs. S4-9), while R is underpredicted by about half of the methods. This can likely be explained by differences in hydration methods’ handling of formally charged amino acids.

Charge matching between the client and scaffold is also likely to influence partitioning of globular clients into condensates. Both scaffold molecules carry a net positive charge, and thus we would expect there to be a preference for negatively-charged clients to partition into the condensate. In our data, we observe that positively charged lysine (K) variant is more excluded from the condensate than the negatively charged glutamate (E) variant, despite glutamate’s high solubility. We also note that E ranks lower than K in underpredictedness for all hydration free energy methods testsed (Figs. S4-9). This supports the notion the charge-matching is an important factor of condensate partitioning that is not captured by calculations of hydration free energy. While Arginine also carries a positive charge, and should thus have some tendency to be excluded from the Nup116 and MacNup98A droplets, we find that it tends to be underpredicted in most cases, but that is likely due to specific interactions with aromatic residues and sp^2^-hybridized groups in the condensate.

Another set of amino acids that are poorly predicted in most cases are the polar uncharged amino acids Q, T, S and N. One possible explanation for this is highlighted in previous work that finds that small-molecule partitioning into condensates is most well-predicted by oil-water transfer[10]. This indicates that the condensate interior behaves more hydrophobic than expected for a predominantly aqueous environment, which may contribute to the overprediction of globular clients decorated with polar uncharged amino acids. Since these residues are soluble in water, their respective GFP variants are already predicted to have a relatively low partitioning into the condensates. However, their partitioning overpredicted, indicating that their presence inside the condensate is even less favorable than predicted by simply just hydration free energy, and the reduced water content inside the condensate. This could be evidence that water inside the condensate behaves differently than bulk water, and is a more favorable environment for hydrophobic molecules, rather than polar molecules. This suggests that the microenvironment inside protein-rich condensates can deviate substantially from bulk solvent properties, though this remains a speculative explanation.

Further insight is gained by comparing the ranking order of GFP variants across experimental data and computational models (Fig. S9). The W variant is consistently ranked as the most underpredicted variant, while the aromatic-rich and arginine-containing variants (R, Y, F) are at least 50% among those with the greatest degree of underpredictedness. In contrast, the polar variants (N, Q, S, T) are consistently ranked lowest, or most overpredicted. Charged variants display more variable behavior across models, reflecting the differing abilities of computational models to capture charge effects. These observations underscore both the strengths and limitations of hydration-based models and emphasize the need for further refinement to account for specific interactions and environmental effects. Future studies should directly incoporate these additoional effects of charge matching, specific interactions, and altered solvent environment to more accurately and robustly predict client partitioning into condensates, and will require a more diverse experimental data set.

## Conclusion

Our results demonstrate that hydration free energy is a strong predictor of the partitioning behavior of GFP variants into FG nucleoporin condensates. Across 17 GFP variants, we observed a robust logarithmic correlation between computed hydration free energy and experimentally measured partition coefficients, supporting the use of hydration free energy as a proxy for transfer free energy in these systems. Variants that are underpredicted by the model are enriched in amino acids capable of specific interactions with aromatic residues of the FG nucleoporins, such as tyrosine, tryptophan, and arginine. This underprediction is likely due to *π*–*π* and cation–*π* interactions that are not fully represented by Δ*F_hyd_*calculations alone. Polar variants are generally overpredicted, suggesting that the condensate interior may be less accommodating to polar side chains due to reduced polarity or solvation capacity. Among the computational methods evaluated, the hi-patch model provides the best agreement with experimental data, indicating that detailed analysis of surface features is critical for accurate prediction of partitioning behavior.

This work provides the ability to predict scaffold-client phyase separation and partitioning behaviors of proteins into condensates, providing quick estimates for different globular proteins and their general partitioning capability into condensates. This predictive capability is valuable for researchers studying phase separation, as it enables the rational selection or design of client proteins for condensate systems prior to carrying out rigorous experimental measurements. These findings inform experimental strategies, facilitate the interpretation of condensate composition, and support the development of novel approaches for assessing the molecular determinants of phase behavior in diverse cellular environments.

## Methods

For each GFP protein variant, eight residues on the surface of the protein were mutated to the same amino acid. For example, GFP variant 8A contains eight surface mutations, all of which were converted to the amino acid Alanine. The same eight regions of the amino acid sequence were modified on each of the seventeen GFP variants, according to the model system variants. The sequences were obtained from Frey et al.[20], and are listed in Table S1. To generate structures of the 17 designed GFP sequences, we used Alphafold3 to generate pdb structures, which were then energy-minimized[58]. We directly used these minimized structures for static ΔF*_tr_* calculations and as starting points for dynamic methods.

The effective energy function 1 (EEF1) of CHARMM was used as it calculates the ΔF*_hyd_* of all atoms without considering the surface area [42]. The adaptive Poisson-Boltzmann solver (APBS) was used because it solved the Poisson Boltzmann equation of the protein and vacuum to approximate the change in ΔF*_hyd_* [49]. The hydrophobic intensity patch (hi-patch) method was adapted to report the hydration free energy. The original implementation reports only hydrophobicity of hydrophobic patches on the surface of the protein, while we report hydration of the full protein surface. The calculations are based off of the EEF1 implicit solvent model[42], and corrected for effects of neighboring atoms, highlighting the cooperative effects of nonpolar atoms in creating hydrophobic interfaces with water[44, 45, 59].

MD simulations were performed with OpenMM in both implicit and explicit solvent setups [60]. We utilized Amber ff14SB for all simulations[51]. Implicit solvent simulations were conducted in the NVT ensemble. Explicit solvent simulations were initially equilibrated in the NVT ensemble, followed by production runs in the NPT ensemble. Temperature was maintained at 300K using the Langevin Middle integrator, and Pressure was kept constant in explicit simulations using a Monte Carlo Barostat. All simulations were conducted at 0.1 M salt concentration and were carried out for 100 ns simulation time. The first 20 ns were discarded for equilibration time.

## Author Contributions

G.L.D. designed the research. S.A. and M.H. performed simulations and analyzed the data. S.A., M.H. and G.L.D. wrote the article.

## Acknowledgments

The authors would like to thank Ben Schuster, José Villegas, Emiliano Brini, and Pradipta Bandyopadhyay for helpful conversations. We also thank Héctor Sánchez-Morán and Joel Kaar for useful conversations and assistance with the hi-patch code. G.L.D. acknowledges support from NIH NIGMS Award number R35GM150589, and start-up funds from Rutgers University. S.A. was supported through the Advanced Materials REU summer fellowship at Rutgers University.

## Declaration of generative AI and AI-assisted technologies in the writing process

During the preparation of this work the authors used Perplexity in order to draft one short section of the paper. After using this tool/service, the authors reviewed and edited the content as needed and take full responsibility for the content of the publication.

## Supplementary Material

An online supplement to this article can be found online. It includes additional information on additional parameters calculated by hi-patch and their predictiveness of partition coefficients. It also includes data from all five additional hydration free energy calculation methods, and their predictiveness of partition coefficients. The tables include descriptions of hi-patch parameters, and amino acid sequences for all proteins used in this work.

## 1. Supplementary Figures

**Fig S1:**
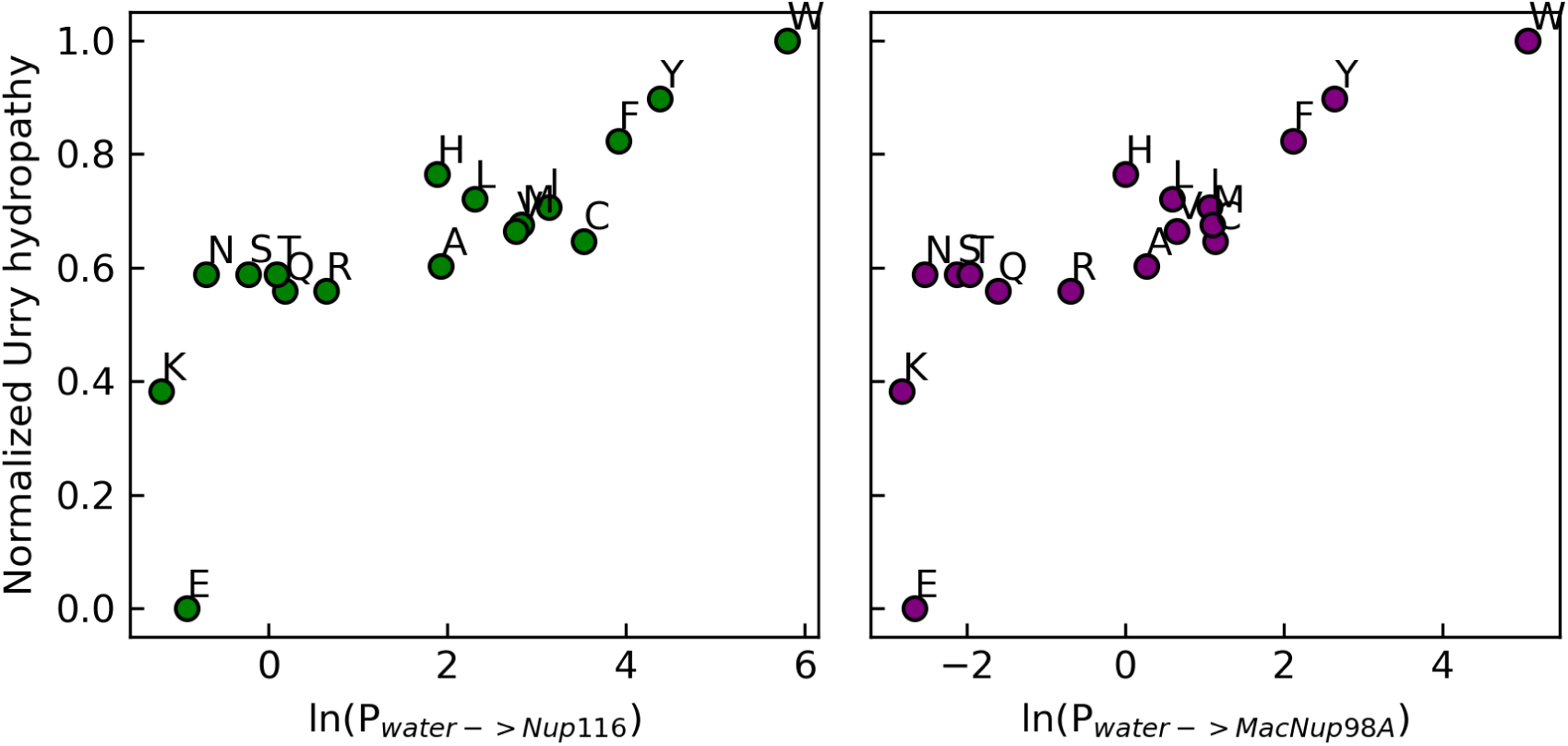
Comparison of experimental partition coefficient^1^ with hydropathy parameters from Urry scale^2^, showing a logarithmic relationship.

**Fig S2:**
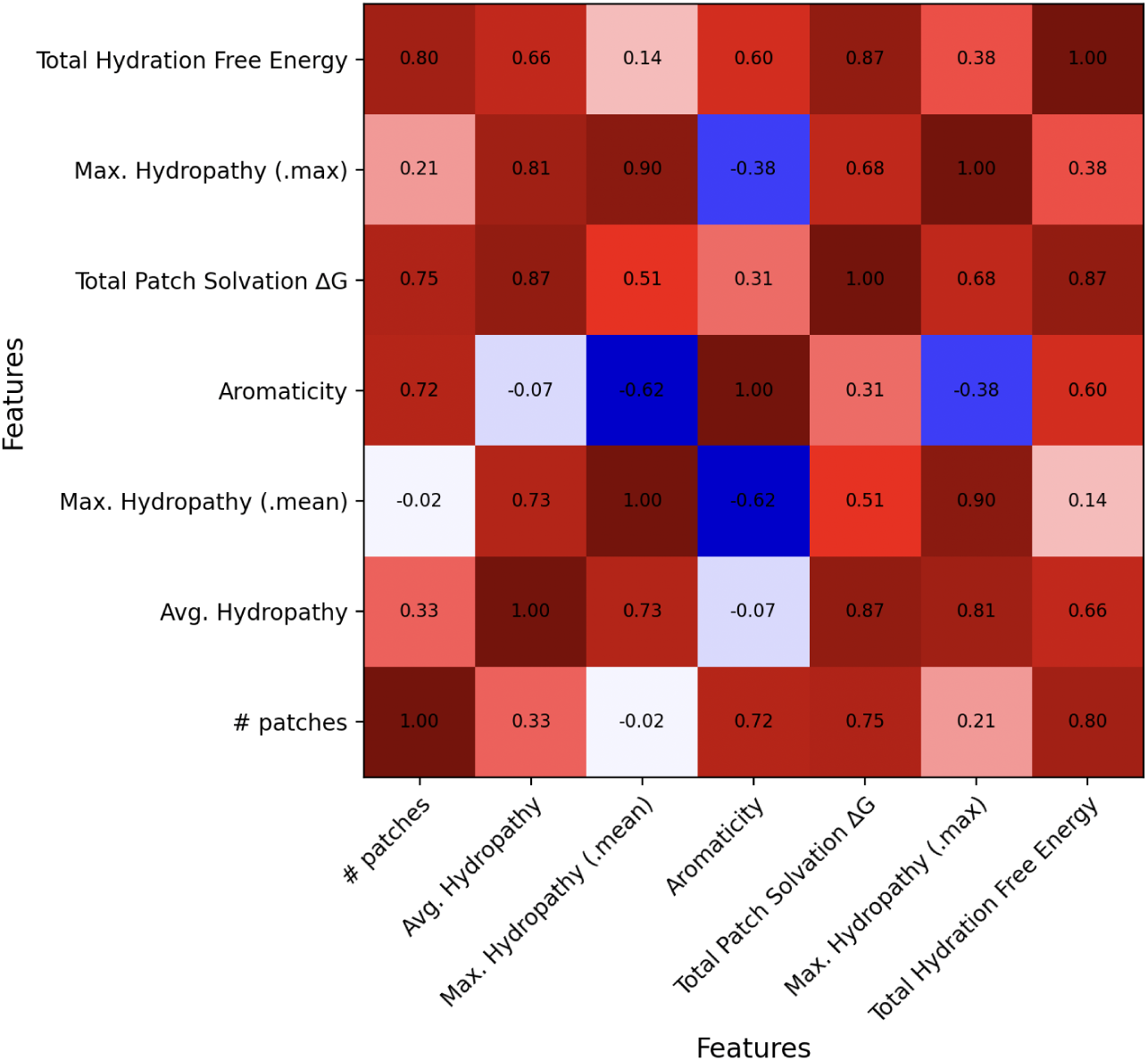
Analysis of different outputs from hi-patch software^3^ (See Table S1) and their relatedness when applied to the 17 variants of GFP. Covariance will be 1 if parameters are perfectly correlated, and -1 if perfectly anticorrelated. A value close to 0 indicates they are uncorrelated.

**Fig. S3:**
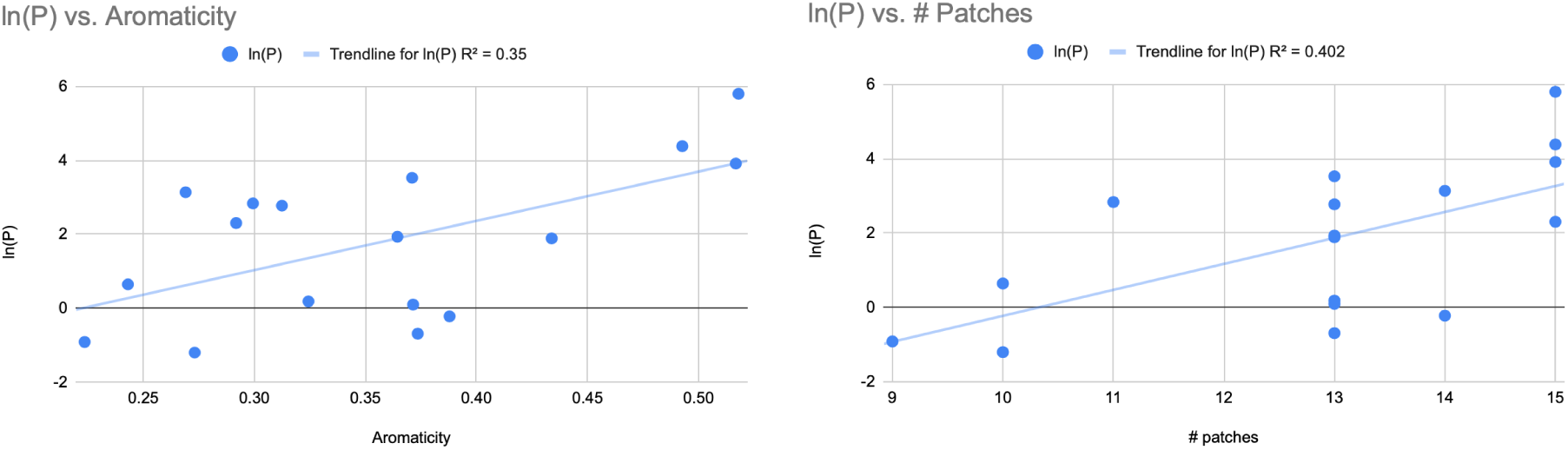
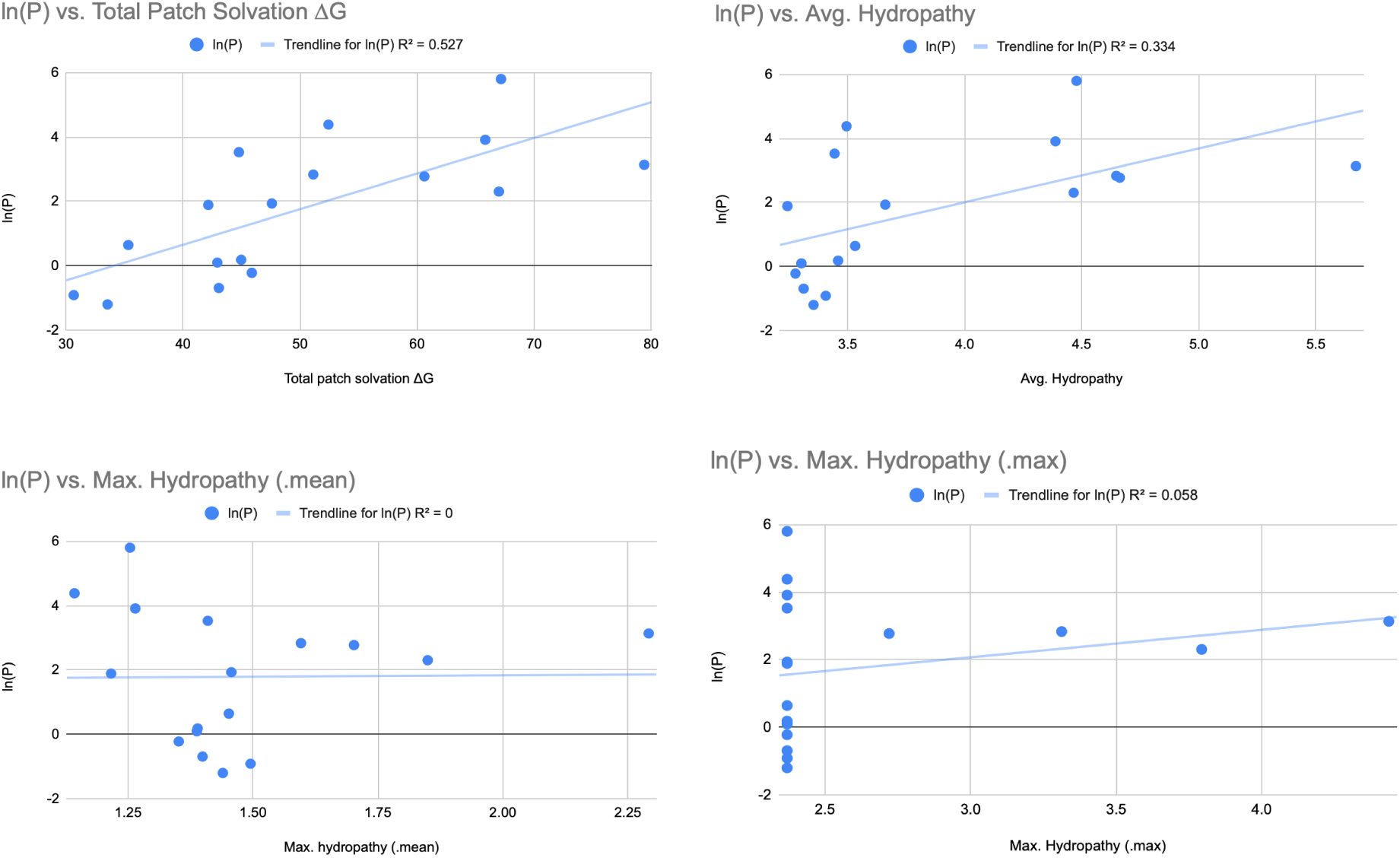
predictions of ln(P) from other hi-patch outputs are not as robust as with the full hydration free energy calculation.

**Fig. S4:**
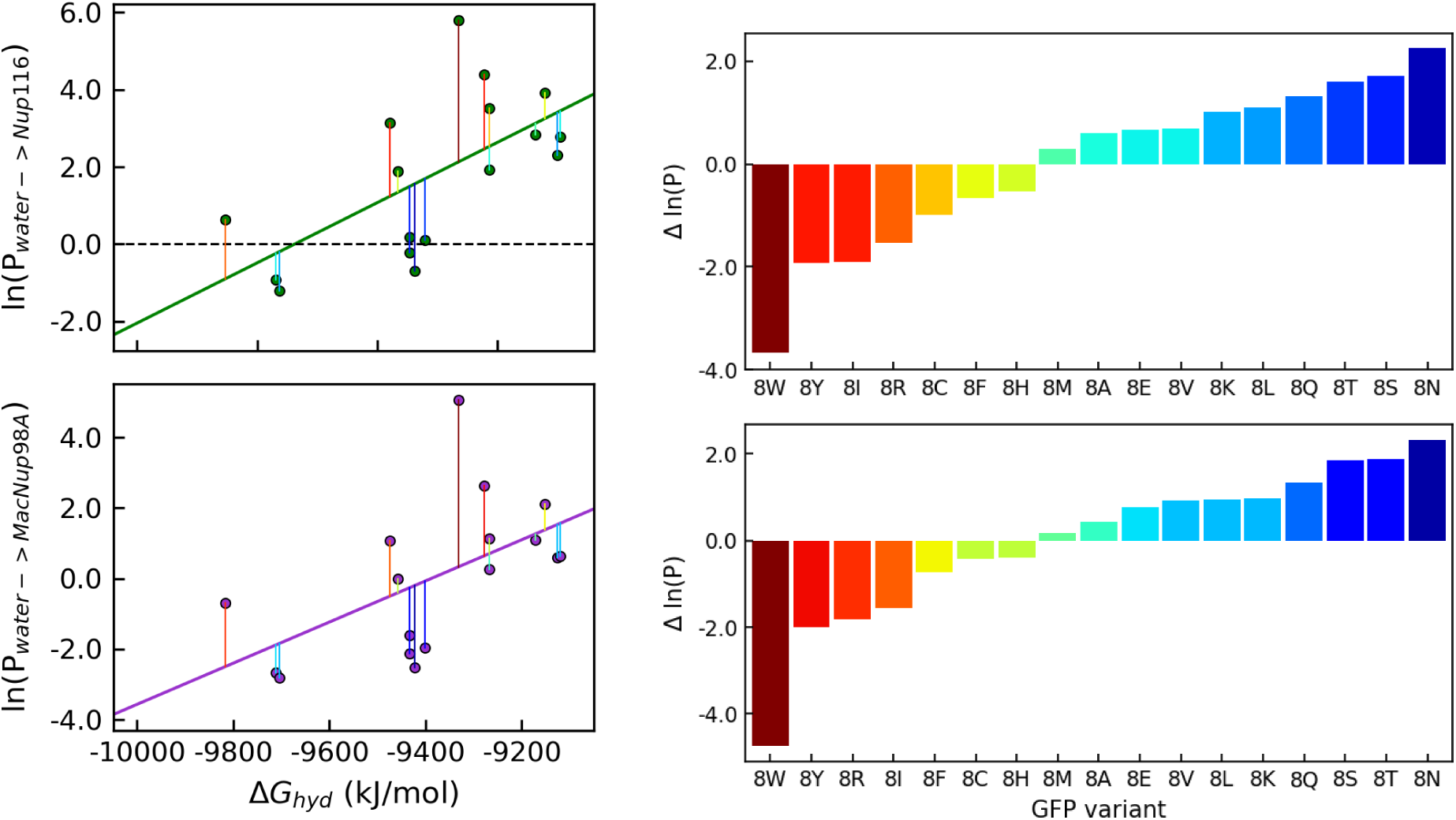
Predictive model and residuals for static calculation of hydration free energy using the eef1 force field implemented in CHARMM^4^.

**Fig S5:**
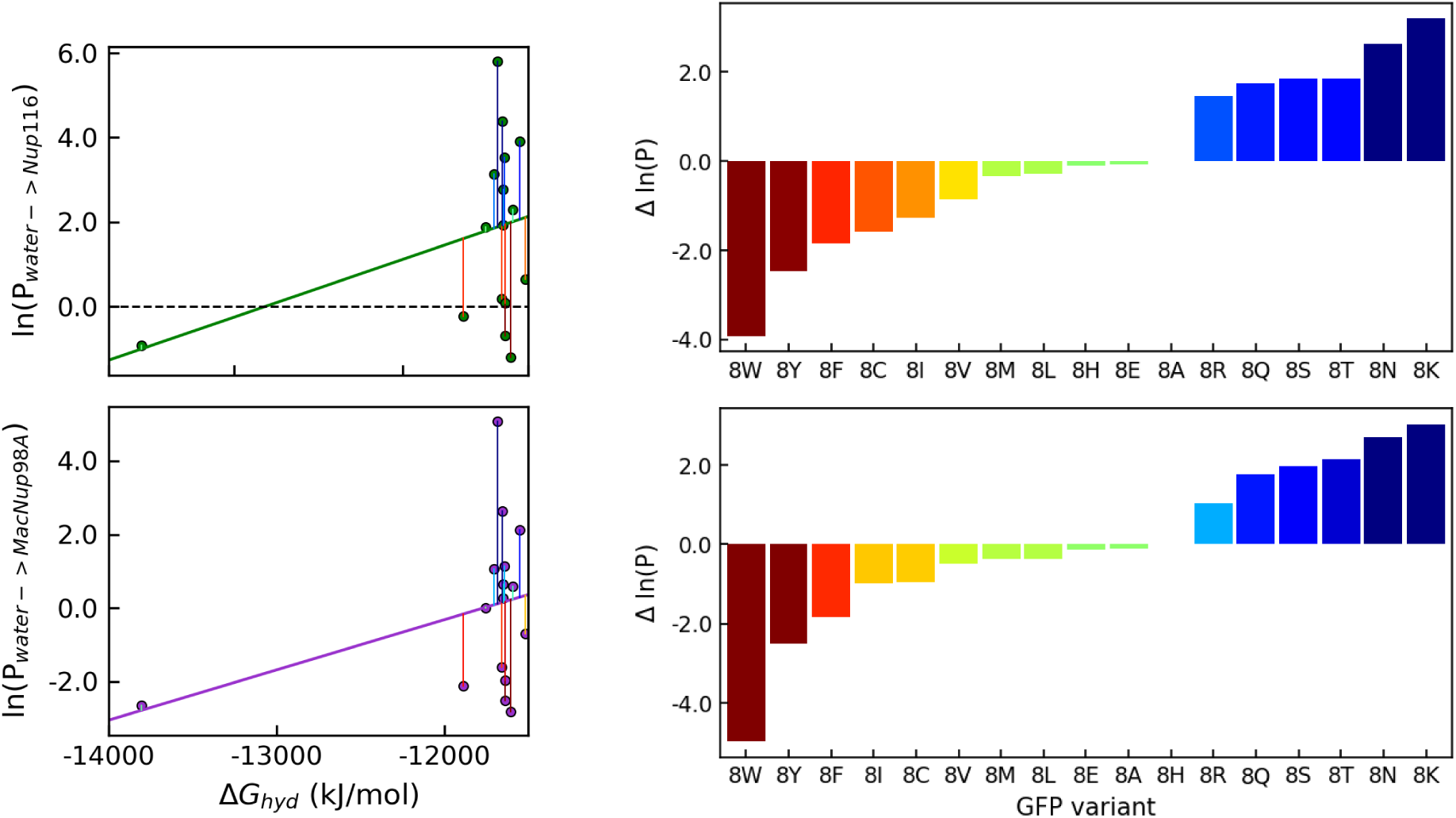
Predictive model and residuals for static calculation of hydration free energy using the adaptive Poisson-Boltzmann solver (APBS)^5^.

**Fig. S6:**
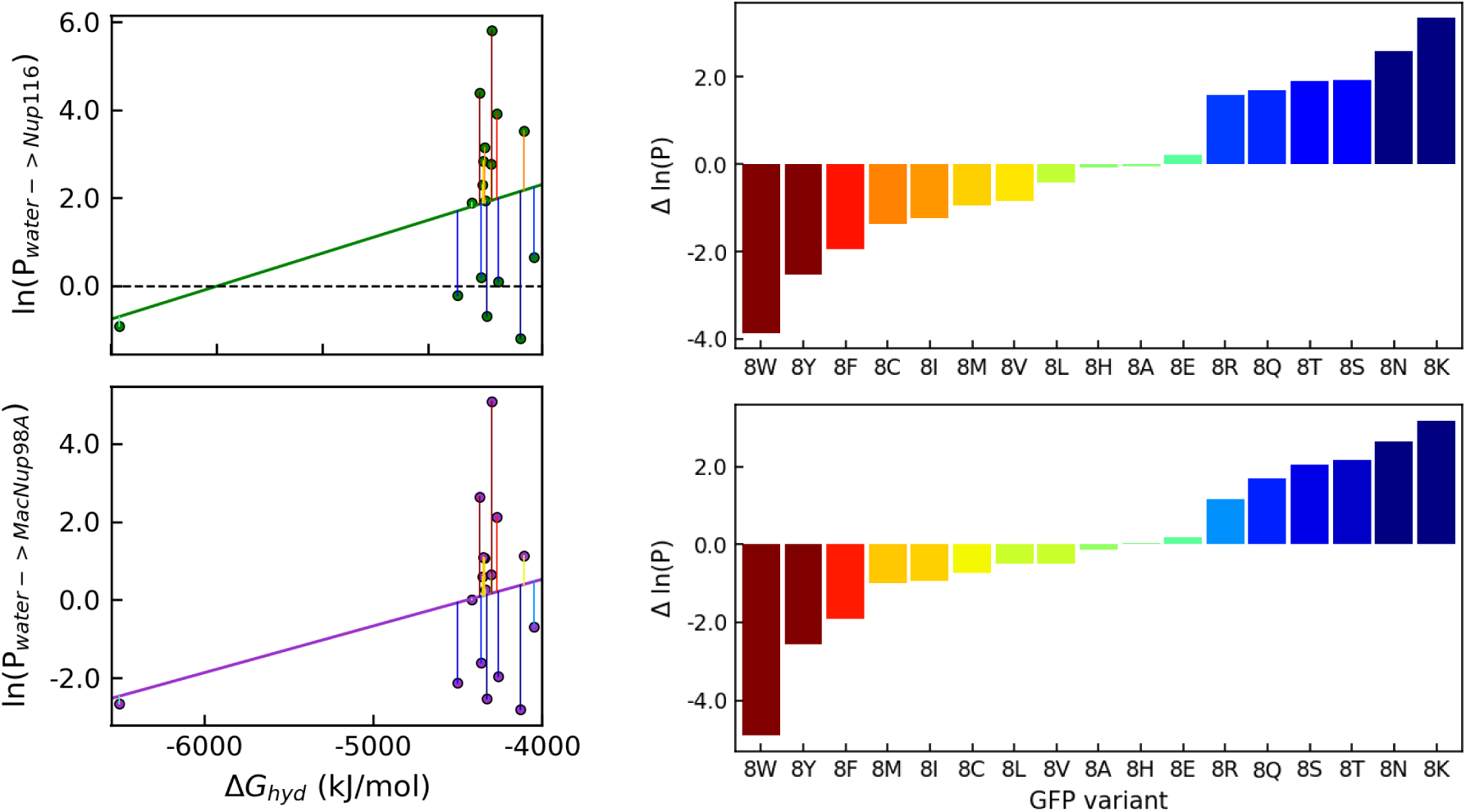
Predictive model and residuals for dynamic hydration free energy calculations using MD simulation with GBNeck implicit solvent^6^.

**Fig. S7:**
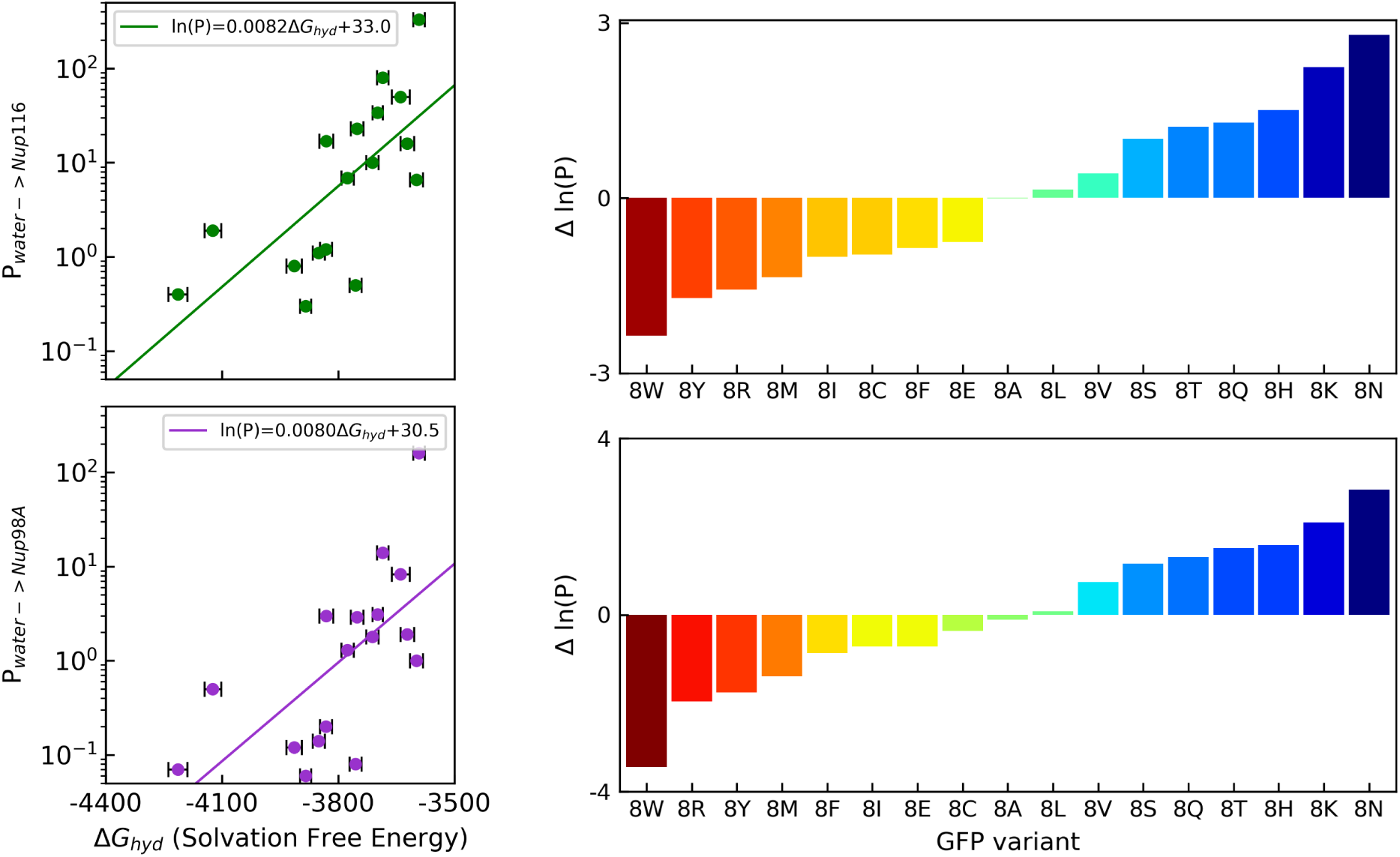
Predictive model and residuals for dynamic explicit solvent simulation trajectory analyzed using hi-patch to calculate hydration free energy. Error bars are SEM from the simulation calculated by block averaging with 5 blocks.

**Figure S8:**
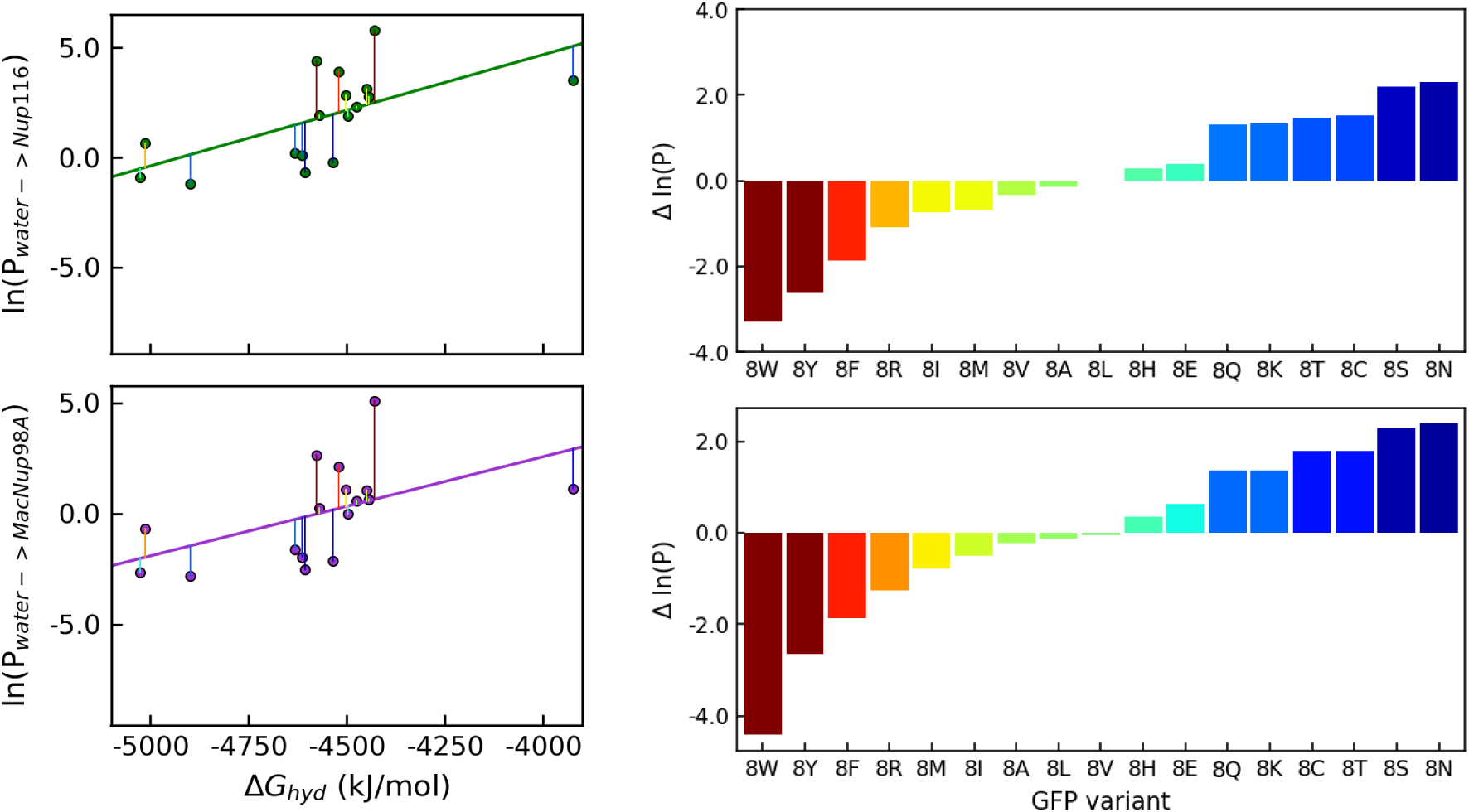
Predictive model and residuals for dynamic implicit solvent simulation trajectory analyzed by hi-patch to calculate hydration free energy.

**Fig. S9:**
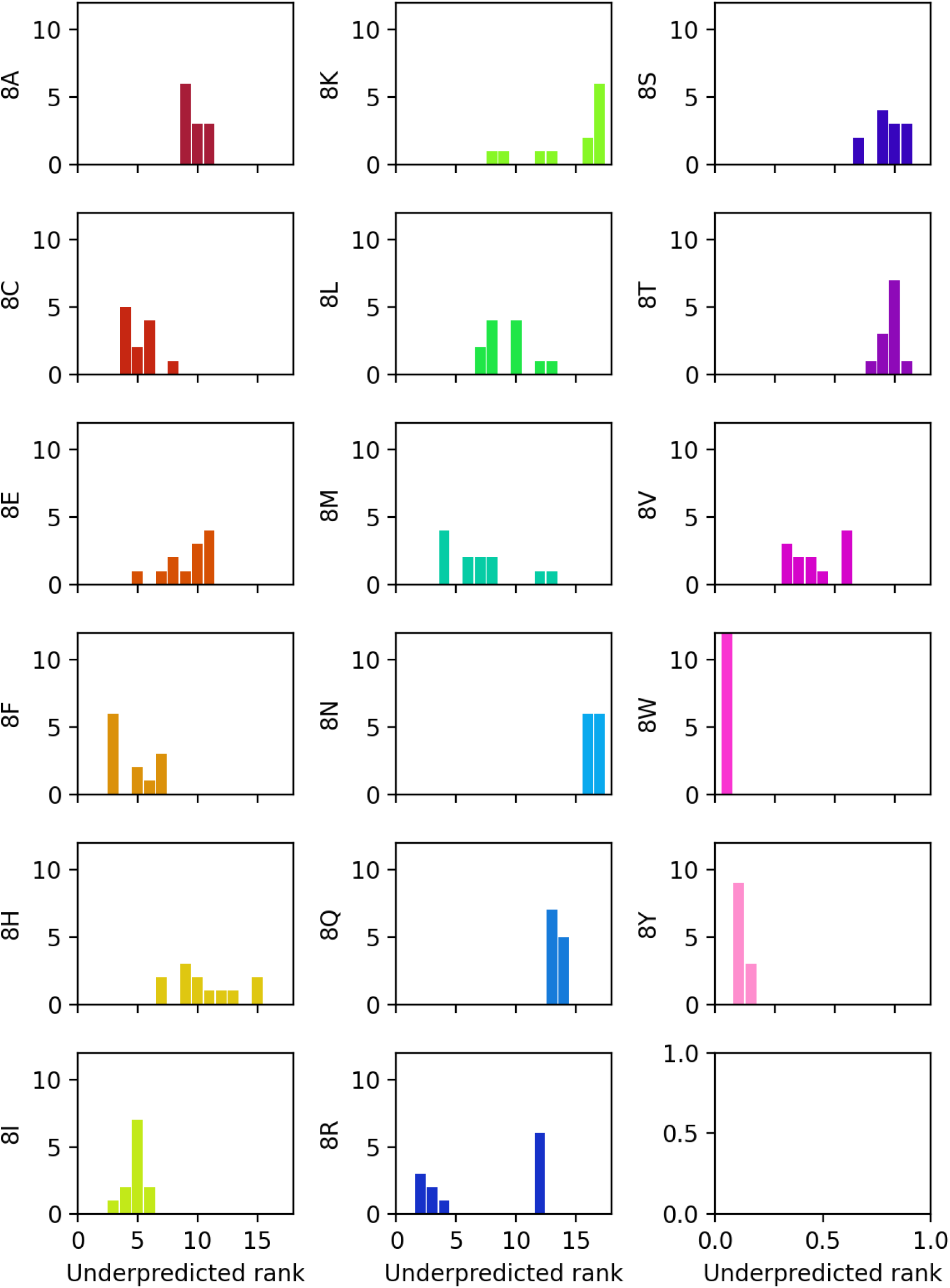
For all six hydration free energy methods tested, we calculated the ranking of overpredicted and underpredicted proteins and show the histogram of ranks for each GFP variant. Having density toward low numbers indicates underpredictedness, and greater specific interactions between the residue type and scaffold proteins

## 2. Supporting Tables

**Table S1:**
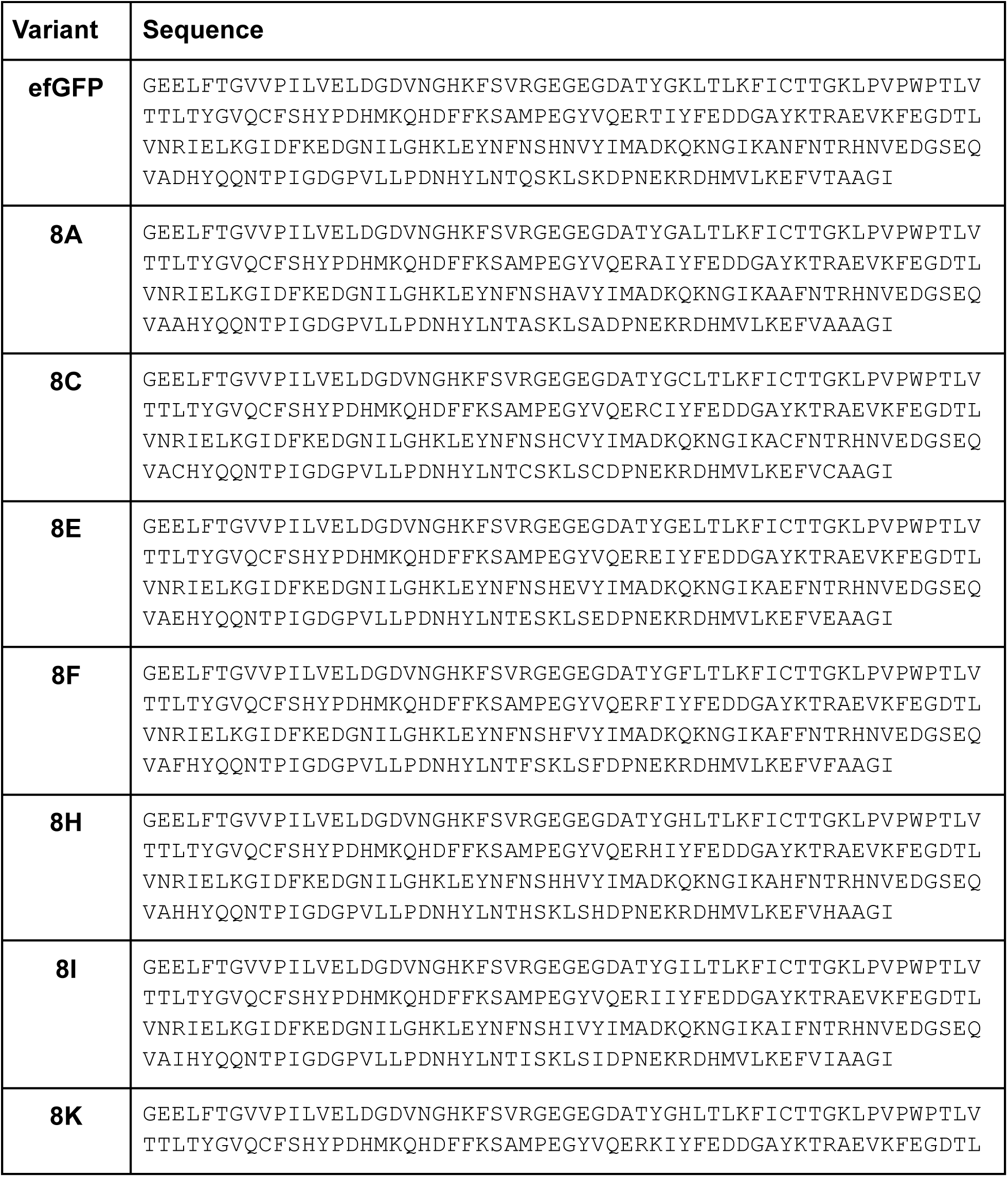
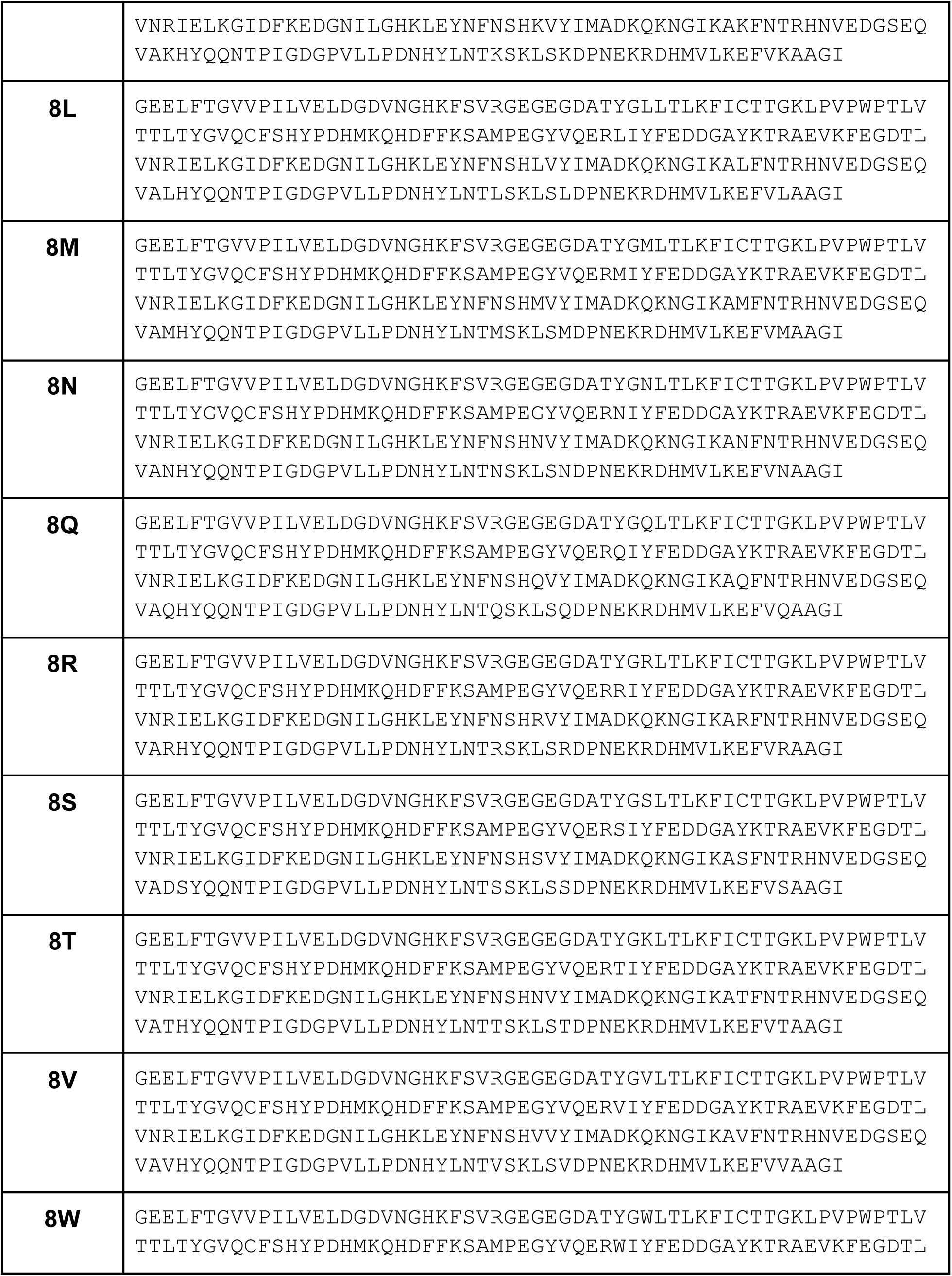
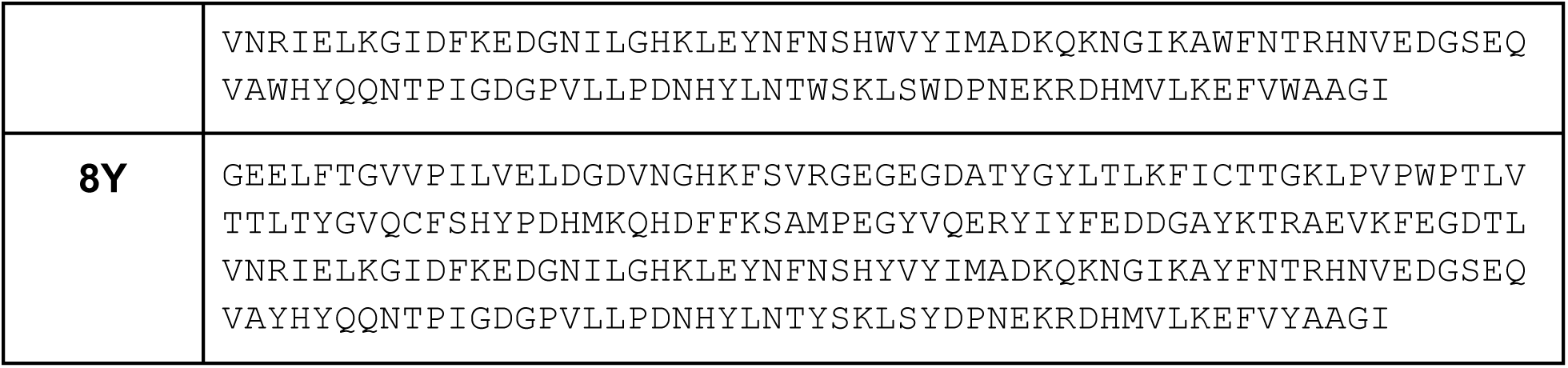
List of GFP variants from Frey et al.^1^ and their amino acid sequences used in modeling of the globular proteins in this work.

**Table S2:**
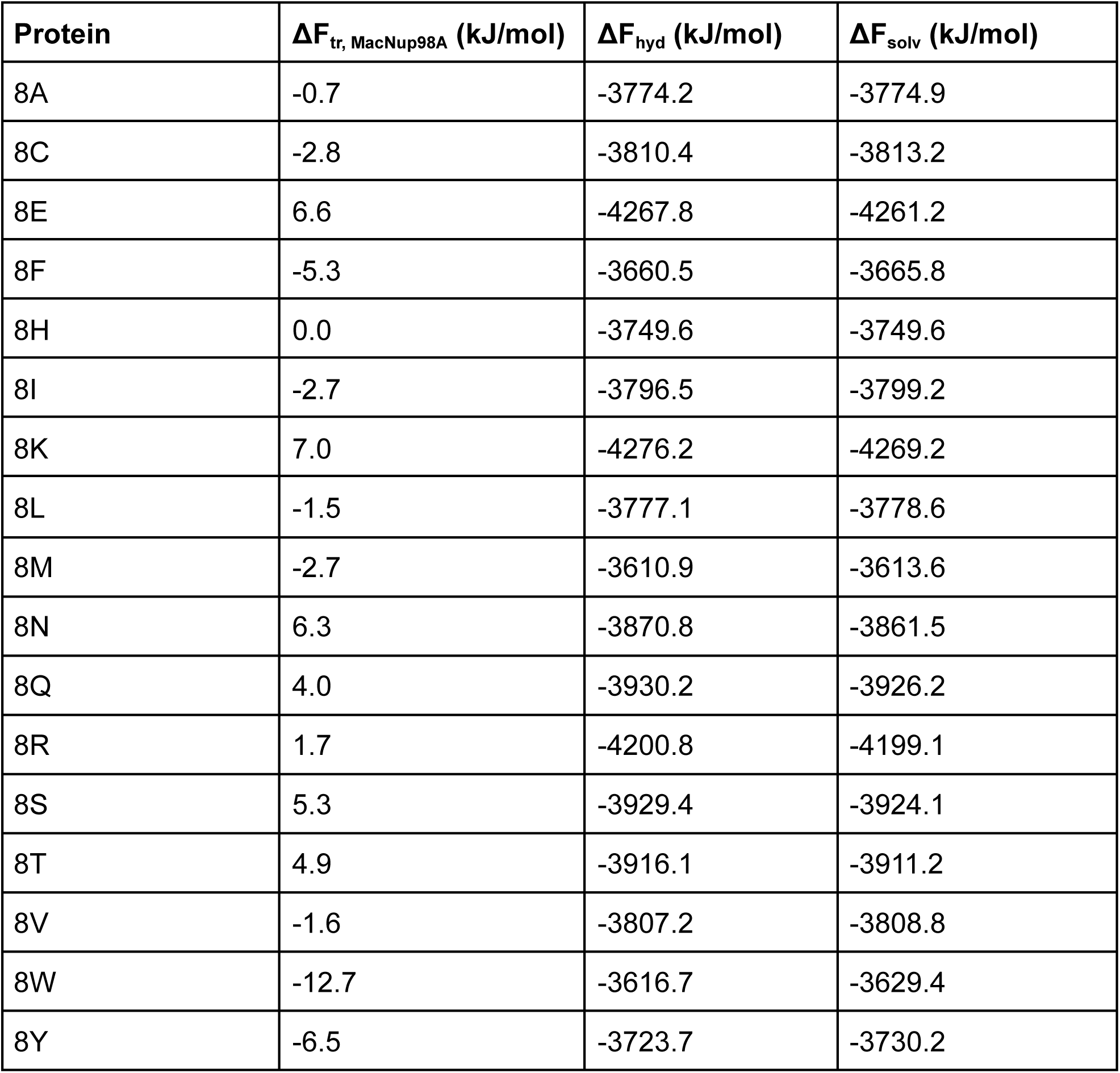
Transfer free energies of each GFP into both scaffold proteins calculated from experiments, and hydration free energies from hi-patch. Solvation free energy is calculated by finding the difference.

| **Protein** | **$\Delta F_{tr, \text{MacNup98A}}$ (kJ/mol)** | **$\Delta F_{hyd}$ (kJ/mol)** | **$\Delta F_{solv}$ (kJ/mol)** |
| --- | --- | --- | --- |
| 8A | -0.7 | -3774.2 | -3774.9 |
| 8C | -2.8 | -3810.4 | -3813.2 |
| 8E | 6.6 | -4267.8 | -4261.2 |
| 8F | -5.3 | -3660.5 | -3665.8 |
| 8H | 0.0 | -3749.6 | -3749.6 |
| 8I | -2.7 | -3796.5 | -3799.2 |
| 8K | 7.0 | -4276.2 | -4269.2 |
| 8L | -1.5 | -3777.1 | -3778.6 |
| 8M | -2.7 | -3610.9 | -3613.6 |
| 8N | 6.3 | -3870.8 | -3861.5 |
| 8Q | 4.0 | -3930.2 | -3926.2 |
| 8R | 1.7 | -4200.8 | -4199.1 |
| 8S | 5.3 | -3929.4 | -3924.1 |
| 8T | 4.9 | -3916.1 | -3911.2 |
| 8V | -1.6 | -3807.2 | -3808.8 |
| 8W | -12.7 | -3616.7 | -3629.4 |
| 8Y | -6.5 | -3723.7 | -3730.2 |

**Table S3:** List and description of parameters calculated by hi-patch software^3^.

|  |  |
| --- | --- |
| # patches | Represents the number of hydrophobic surface regions identified. These patches represent clustered hydrophobic residues exposed to solvent. |
| Avg. Hydropathy | The mean hydropathy value for each protein variant. |
| Max. Hydropathy (.mean) | The highest average hydropathy value among all individual surface patches, representing the most hydrophobic surface region for each variant. |
| Aromaticity | Quantifies the presence of aromatic residues (Tyr, Trp, Phe) on the protein's surface. |
| Total Patch Solvation $\Delta G$ | The total solvation free energy contribution from all identified hydrophobic patches. |
| Max. Hydropathy (.max) | The single highest hydropathy value at any atomic position on the protein surface, identifying the most hydrophobic individual site for each variant. |
| Total Hydration Free Energy | The total free energy change ($\Delta F_{\text{hyd}}$) when transferring a protein from vacuum to aqueous solution. |

**Table S4:**
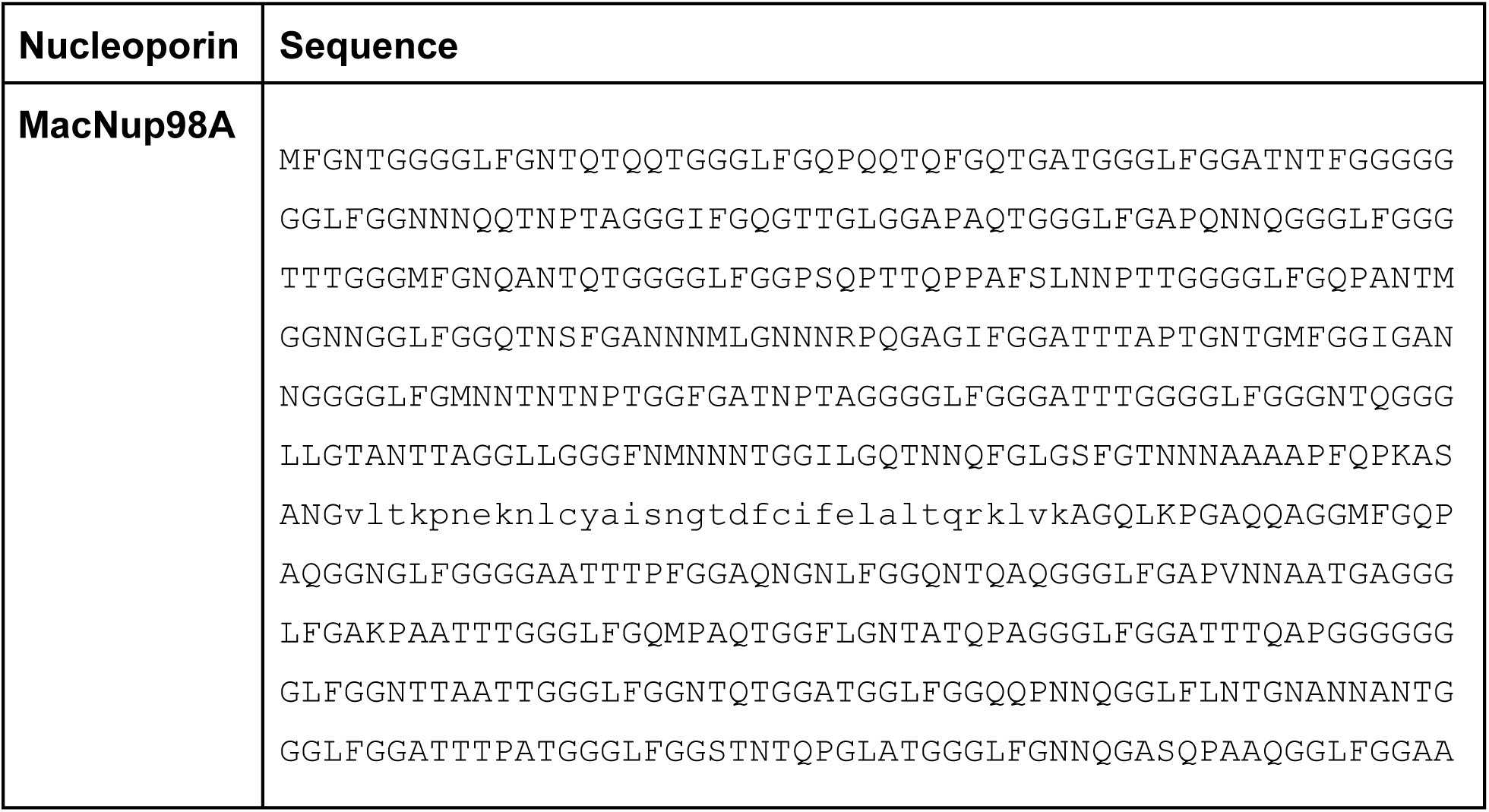
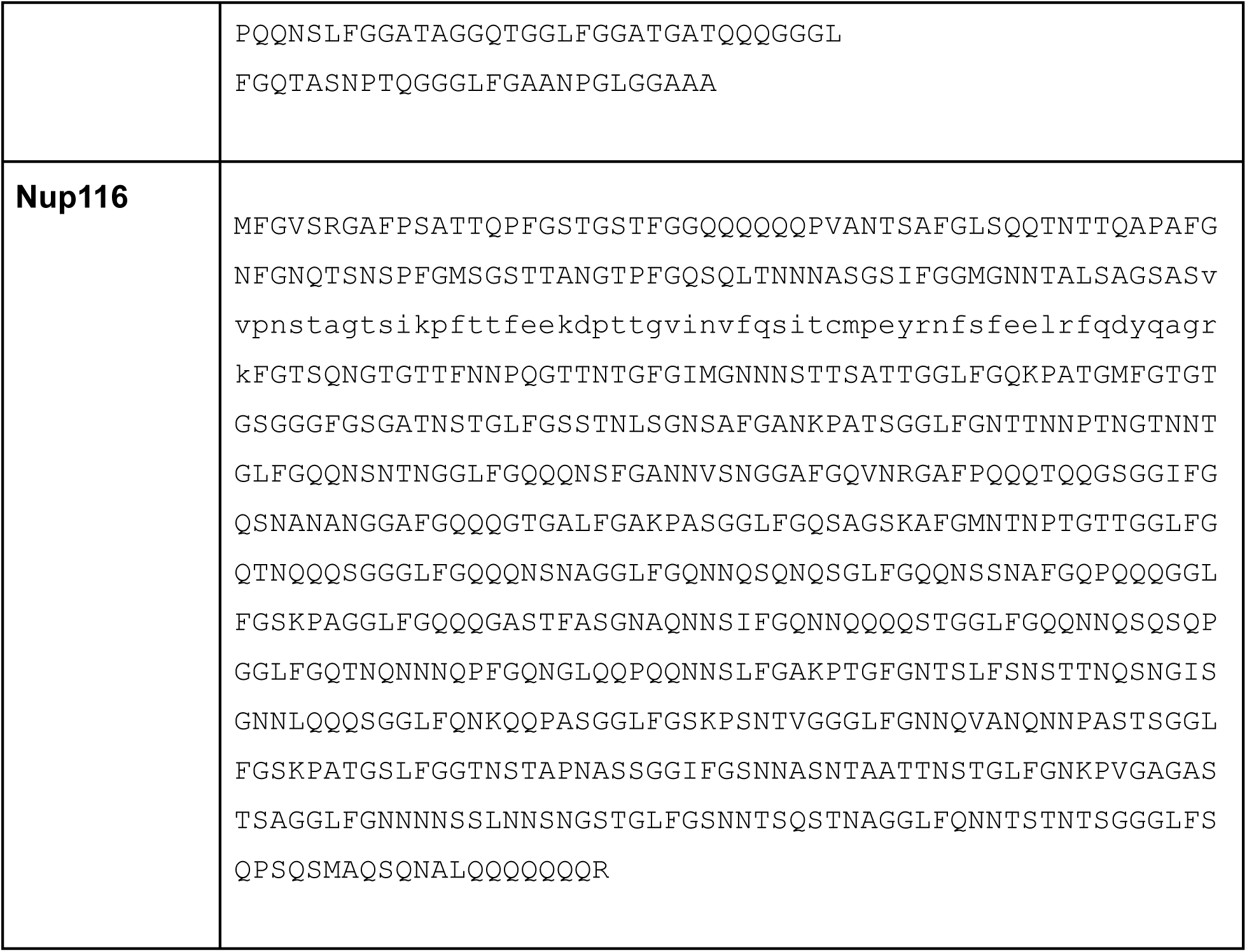
Sequences for the two Nucleoporin proteins from Frey et al.^1^.

**Table S5:** Size, charge and hydropathy^2^ of protein sequences used in this work.

| Name | Length | Net charge | Urry hydropathy |
| --- | --- | --- | --- |
| 8A | 226 | -6 | 0.573 |
| 8C | 226 | -6 | 0.575 |
| 8E | 226 | -14 | 0.552 |
| 8F | 226 | -6 | 0.581 |
| 8H | 226 | -2 | 0.579 |
| 8I | 226 | -6 | 0.577 |
| 8K | 226 | 1.5 | 0.567 |
| 8L | 226 | -6 | 0.577 |
| 8M | 226 | -6 | 0.576 |
| 8N | 226 | -6 | 0.572 |
| 8Q | 226 | -6 | 0.571 |
| 8R | 226 | 2.0 | 0.571 |
| 8S | 226 | -7.5 | 0.570 |
| 8T | 226 | -5 | 0.572 |
| 8V | 226 | -6 | 0.575 |
| 8W | 226 | -6 | 0.587 |
| 8Y | 226 | -6 | 0.583 |
| Nup116 | 1113 | 20.5 | 0.592 |
| MacNup98A | 1105 | 16 | 0.594 |

## References

[1] Christine D Keating. Aqueous phase separation as a possible route to compartmentalization of biological molecules. Accounts of chemical research, 45(12):2114–2124, 2012.

[2] Yongdae Shin and Clifford P Brangwynne. Liquid phase condensation in cell physiology and disease. Science, 357(6357):eaaf4382, 2017.

[3] Salman F. Banani, Hyun O. Lee, Anthony A. Hyman, and Michael K. Rosen. Biomolecular condensates: Organizers of cellular biochemistry. Nature Reviews Molecular Cell Biology, 18(5):285–298, 2017. Publisher: Nature Publishing Group.

[4] Gregory L Dignon, Robert B Best, and Jeetain Mittal. Biomolecular phase separation: from molecular driving forces to macroscopic properties. Annual review of physical chemistry, 71:53–75, 2020.

[5] William M. Jacobs and Daan Frenkel. Phase Transitions in Biological Systems with Many Components. Biophysical Journal, 112(4):683–691, 2017. arXiv: 1703.01223 Publisher: Biophysical Society ISBN: 0006-3495.

[6] Shana Elbaum-Garfinkle, Younghoon Kim, Krzysztof Szczepaniak, Carlos Chih-Hsiung Chen, Christian R Eckmann, Sua Myong, and Clifford P Brangwynne. The disordered p granule protein laf-1 drives phase separation into droplets with tunable viscosity and dynamics. Proceedings of the National Academy of Sciences, 112(23):7189–7194, 2015.

[7] Kathleen A Burke, Abigail M Janke, Christy L Rhine, and Nicolas L Fawzi. Residue-by-residue view of in vitro fus granules that bind the c-terminal domain of rna polymerase ii. Molecular cell, 60(2):231–241, 2015.

[8] Julie D Forman-Kay, Richard W Kriwacki, and Geraldine Seydoux. Phase separation in biology and disease. Journal of molecular biology, 430(23):4603, 2018.

[9] Huan Wang, Fleurie M. Kelley, Dragomir Milovanovic, Benjamin S. Schuster, and Zheng Shi. Surface tension and viscosity of protein condensates quantified by micropipette aspiration. Biophysical Reports, 1(1):100011, September 2021.

[10] Sabareesan Ambadi Thody, Hanna D. Clements, Hamid Baniasadi, Andrew S. Lyon, Matthew S. Sigman, and Michael K. Rosen. Small-molecule properties define partitioning into biomolecular condensates. Nature Chemistry, 16(11):1794–1802, November 2024.

[11] P Andrew Chong and Julie D Forman-Kay. Liquid–liquid phase separation in cellular signaling systems. Current opinion in structural biology, 41:180–186, 2016.

[12] Mikael V Garabedian, Wentao Wang, Jorge B Dabdoub, Michelle Tong, Reese M Caldwell, William Benman, Benjamin S Schuster, Alexander Deiters, and Matthew C Good. Designer membraneless organelles sequester native factors for control of cell behavior. Nature chemical biology, 17(9):998–1007, 2021.

[13] Samantha Keyport Kik, Dana Christopher, Hendrik Glauninger, Caitlin Wong Hickernell, Jared AM Bard, Kyle M Lin, Allison H Squires, Michael Ford, Tobin R Sosnick, and D Allan Drummond. An adaptive biomolecular condensation response is conserved across environmentally divergent species. Nature Communications, 15(1):3127, 2024.

[14] Christian Hoffmann, Jakob Rentsch, Taka A Tsunoyama, Akshita Chhabra, Gerard Aguilar Perez, Rajdeep Chowdhury, Franziska Trnka, Aleksandr A Korobeinikov, Ali H Shaib, Marcelo Ganzella, et al. Synapsin condensation controls synaptic vesicle sequestering and dynamics. Nature Communications, 14(1):6730, 2023.

[15] Danfeng Cai, Zhe Liu, and Jennifer Lippincott-Schwartz. Biomolecular condensates and their links to cancer progression. Trends in biochemical sciences, 46(7):535–549, 2021.

[16] Huaiying Zhang, Shana Elbaum-Garfinkle, Erin M Langdon, Nicole Taylor, Patricia Occhipinti, Andrew A Bridges, Clifford P Brangwynne, and Amy S Gladfelter. Rna controls polyq protein phase transitions. Molecular cell, 60(2):220–230, 2015.

[17] Yuan Lin, David SW Protter, Michael K Rosen, and Roy Parker. Formation and maturation of phase-separated liquid droplets by rna-binding proteins. Molecular cell, 60(2):208–219, 2015.

[18] Avinash Patel, Hyun O Lee, Louise Jawerth, Shovamayee Maharana, Marcus Jahnel, Marco Y Hein, Stoyno Stoynov, Julia Mahamid, Shambaditya Saha, Titus M Franzmann, et al. A liquid-to-solid phase transition of the als protein fus accelerated by disease mutation. Cell, 162(5):1066–1077, 2015.

[19] Salman F Banani, Allyson M Rice, William B Peeples, Yuan Lin, Saumya Jain, Roy Parker, and Michael K Rosen. Compositional control of phase-separated cellular bodies. Cell, 166(3):651–663, 2016.

[20] Steffen Frey, Renate Rees, Jürgen Schünemann, Sheung Chun Ng, Kevser Fünfgeld, Trevor Huyton, and Dirk Görlich. Surface properties determining passage rates of proteins through nuclear pores. Cell, 174(1):202–217, 2018.

[21] Wenmin Xing, Denise Muhlrad, Roy Parker, and Michael K Rosen. A quantitative inventory of yeast p body proteins reveals principles of composition and specificity. Elife, 9:e56525, 2020.

[22] Fleurie M Kelley, Anas Ani, Emily G Pinlac, Bridget Linders, Bruna Favetta, Mayur Barai, Yuchen Ma, Arjun Singh, Gregory L Dignon, Yuwei Gu, et al. Controlled and orthogonal partitioning of large particles into biomolecular condensates. Nature Communications, 16(1):3521, 2025.

[23] Patrick M McCall, Kyoohyun Kim, Anatol W Fritsch, JM Iglesias-Artola, LM Jawerth, Jie Wang, Martine Ruer, J Peychl, Andrey Poznyakovskiy, Jochen Guck, et al. Quantitative phase microscopy enables precise and efficient determination of biomolecular condensate composition. BioRxiv, pages 2020–10, 2020.

[24] Daoyuan Qian, Hannes Ausserwoger, Tomas Sneideris, Mina Farag, Rohit V Pappu, and Tuomas PJ Knowles. Dominance analysis to assess solute contributions to multicomponent phase equilibria. Proceedings of the National Academy of Sciences, 121(33):e2407453121, 2024.

[25] Yi Hsuan Lin, Jacob P. Brady, Julie D. Forman-Kay, and Hue Sun Chan. Charge pattern matching as a ’fuzzy’ mode of molecular recognition for the functional phase separations of intrinsically disordered proteins. New Journal of Physics, 19(11):115003, 2017.

[26] Jie Wang, Jeong-Mo Choi, Alex S Holehouse, Hyun O Lee, Xiaojie Zhang, Marcus Jahnel, Shovamayee Maharana, Régis Lemaitre, Andrei Pozniakovsky, David Drechsel, et al. A molecular grammar governing the driving forces for phase separation of prion-like rna binding proteins. Cell, 174(3):688–699, 2018.

[27] Gregory L Dignon, Wenwei Zheng, Young C Kim, Robert B Best, and Jeetain Mittal. Sequence determinants of protein phase behavior from a coarse-grained model. PLoS computational biology, 14(1):e1005941, 2018.

[28] Shiv Rekhi, Cristobal Garcia Garcia, Mayur Barai, Azamat Rizuan, Benjamin S Schuster, Kristi L Kiick, and Jeetain Mittal. Expanding the molecular language of protein liquid–liquid phase separation. Nature Chemistry, pages 1–12, 2024.

[29] Anastasia C Murthy, Gregory L Dignon, Yelena Kan, Gül H Zerze, Sapun H Parekh, Jeetain Mittal, and Nicolas L Fawzi. Molecular interactions underlying liquid-liquid phase separation of the fus low-complexity domain. Nature structural & molecular biology, 26(7):637–648, 2019.

[30] Felipe Garćıa Quiroz and Ashutosh Chilkoti. Sequence heuristics to encode phase behaviour in intrinsically disordered protein polymers. Nature Materials, 14(11):1164–1171, 2015.

[31] Chad Gu, Andrei Vovk, Tiantian Zheng, Rob D Coalson, and Anton Zilman. The role of cohesiveness in the permeability of the spatial assemblies of fg nucleoporins. Biophysical journal, 116(7):1204–1215, 2019.

[32] Joshua A Riback, Micayla A Bowman, Adam M Zmyslowski, Catherine R Knoverek, John M Jumper, James R Hinshaw, Emily B Kaye, Karl F Freed, Patricia L Clark, and Tobin R Sosnick. Innovative scattering analysis shows that hydrophobic disordered proteins are expanded in water. Science, 358(63601):238–241, 2017.

[33] Vivian Yeong, Jou-wen Wang, Justin M Horn, and Allie C Obermeyer. Intracellular phase separation of globular proteins facilitated by short cationic peptides. Nature Communications, 13(1):7882, 2022.

[34] Hang Kuen Lau, Linqing Li, Anna K Jurusik, Chandran R Sabanayagam, and Kristi L Kiick. Aqueous liquid–liquid phase separation of resilin-like polypeptide/polyethylene glycol solutions for the formation of microstructured hydrogels. ACS Biomaterials Science & Engineering, 3(5):757–766, 2017.

[35] So Yeon Ahn and Allie C Obermeyer. Selectivity of complex coacervation in multiprotein mixtures. JACS Au, 4(10):3800–3812, 2024.

[36] Marina Feric, Nilesh Vaidya, Tyler S Harmon, Diana M Mitrea, Lian Zhu, Tiffany M Richardson, Richard W Kriwacki, Rohit V Pappu, and Clifford P Brangwynne. Coexisting liquid phases underlie nucleolar subcompartments. Cell, 165(7):1686–1697, 2016.

[37] Richard C Remsing, Erte Xi, and Amish J Patel. Protein hydration thermodynamics: The influence of flexibility and salt on hydrophobin ii hydration. The Journal of Physical Chemistry B, 122(13):3635– 3646, 2018.

[38] Christopher J Fennell, Charlie Kehoe, and Ken A Dill. Oil/water transfer is partly driven by molecular shape, not just size. Journal of the American Chemical Society, 132(1):234–240, 2010.

[39] Joshua A Riback, Lian Zhu, Mylene C Ferrolino, Michele Tolbert, Diana M Mitrea, David W Sanders, Ming-Tzo Wei, Richard W Kriwacki, and Clifford P Brangwynne. Composition-dependent thermodynamics of intracellular phase separation. Nature, 581(7807):209–214, 2020.

[40] Dan W. Urry, D. Channe Gowda, Timothy M. Parker, Chi-Hao -H Luan, Michael C. Reid, Cynthia M. Harris, Asima Pattanaik, and R. Dean Harris. Hydrophobicity scale for proteins based on inverse temperature transitions. Biopolymers, 32(9):1243–1250, 1992. ISBN: 0006-3525.

[41] Roshan Mammen Regy, Jacob Thompson, Young C. Kim, and Jeetain Mittal. Improved coarse-grained model for studying sequence dependent phase separation of disordered proteins. Protein Science, 30(7):1371–1379, 2021. eprint: https://onlinelibrary.wiley.com/doi/pdf/10.1002/pro.4094.

[42] Themis Lazaridis and Martin Karplus. Effective energy function for proteins in solution. Proteins: Structure, Function, and Bioinformatics, 35(2):133–152, 1999.

[43] Daniel K. Schwartz Héctor Sánchez-Morán James S. Weltz and Joel L. Kaar. Understanding design rules for optimizing the interface between immobilized enzymes and random copolymer brushes. ACS Applied Materials and Interfaces, 2021.

[44] Nicholas B Rego, Erte Xi, and Amish J Patel. Identifying hydrophobic protein patches to inform protein interaction interfaces. Proceedings of the National Academy of Sciences, 118(6):e2018234118, 2021.

[45] Nicholas B Rego, Andrew L Ferguson, and Amish J Patel. Learning the relationship between nanoscale chemical patterning and hydrophobicity. Proceedings of the National Academy of Sciences, 119(48):e2200018119, 2022.

[46] Erik W. Martin, Alex S. Holehouse, Ivan Peran, Mina Farag, J. Jeremias Incicco, Anne Bremer, Christy R. Grace, Andrea Soranno, Rohit V. Pappu, and Tanja Mittag. Valence and patterning of aromatic residues determine the phase behavior of prion-like domains. Science, 367(6478):694–699, 2020.

[47] Henry R Kilgore, Peter G Mikhael, Kalon J Overholt, Ann Boija, Nancy M Hannett, Catherine Van Dongen, Tong Ihn Lee, Young-Tae Chang, Regina Barzilay, and Richard A Young. Distinct chemical environments in biomolecular condensates. Nature Chemical Biology, 20(3):291–301, 2024.

[48] Bernard R Brooks, Charles L Brooks III, Alexander D Mackerell Jr, Lennart Nilsson, Robert J Petrella, Benoît Roux, Youngdo Won, Georgios Archontis, Christian Bartels, Stefan Boresch, et al. Charmm: the biomolecular simulation program. Journal of computational chemistry, 30(10):1545–1614, 2009.

[49] Elizabeth Jurrus, Dave Engel, Keith Star, Kyle Monson, Juan Brandi, Lisa E Felberg, David H Brookes, Leighton Wilson, Jiahui Chen, Karina Liles, et al. Improvements to the apbs biomolecular solvation software suite. Protein science, 27(1):112–128, 2018.

[50] John Mongan, Carlos Simmerling, J Andrew McCammon, David A Case, and Alexey Onufriev. Generalized born model with a simple, robust molecular volume correction. Journal of chemical theory and computation, 3(1):156–169, 2007.

[51] James A Maier, Carmenza Martinez, Koushik Kasavajhala, Lauren Wickstrom, Kevin E Hauser, and Carlos Simmerling. ff14sb: improving the accuracy of protein side chain and backbone parameters from ff99sb. Journal of chemical theory and computation, 11(8):3696–3713, 2015.

[52] Rakesh Srivastava, Mausumi Chattopadhyaya, and Pradipta Bandyopadhyay. Calculation of salt-dependent free energy of binding of *β*-lactoglobulin homodimer formation and mechanism of dimer formation using molecular dynamics simulation and three-dimensional reference interaction site model (3d-rism): diffuse salt ions and non-polar interactions between the monomers favor the dimer formation. Physical Chemistry Chemical Physics, 22(4):2142–2156, 2020.

[53] Shiv Rekhi and Jeetain Mittal. Amino acid transfer free energies reveal thermodynamic driving forces in biomolecular condensate formation. bioRxiv, pages 2024–12, 2024.

[54] Benjamin S Schuster, Gregory L Dignon, Wai Shing Tang, Fleurie M Kelley, Aishwarya Kanchi Ranganath, Craig N Jahnke, Alison G Simpkins, Roshan Mammen Regy, Daniel A Hammer, Matthew C Good, et al. Identifying sequence perturbations to an intrinsically disordered protein that determine its phase-separation behavior. Proceedings of the National Academy of Sciences, 117(21):11421–11431, 2020.

[55] Michael Rubinstein and Andrey V Dobrynin. Solutions of associative polymers. Trends in Polymer Science, 5(6):181–186, 1997.

[56] Jeong-Mo Choi, Alex S Holehouse, and Rohit V Pappu. Physical principles underlying the complex biology of intracellular phase transitions. Annual review of biophysics, 49(1):107–133, 2020.

[57] Robert McCoy Vernon, Paul Andrew Chong, Brian Tsang, Tae Hun Kim, Alaji Bah, Patrick Farber, Hong Lin, and Julie Deborah Forman-Kay. Pi-pi contacts are an overlooked protein feature relevant to phase separation. elife, 7:e31486, 2018.

[58] Josh Abramson, Jonas Adler, Jack Dunger, Richard Evans, Tim Green, Alexander Pritzel, Olaf Ronneberger, Lindsay Willmore, Andrew J Ballard, Joshua Bambrick, et al. Accurate structure prediction of biomolecular interactions with alphafold 3. Nature, 630(8016):493–500, 2024.

[59] Hari Acharya, Srivathsan Vembanur, Sumanth N Jamadagni, and Shekhar Garde. Mapping hydrophobicity at the nanoscale: Applications to heterogeneous surfaces and proteins. Faraday discussions, 146:353–365, 2010.

[60] Peter Eastman, Raimondas Galvelis, Raúl P Peláez, Charlles RA Abreu, Stephen E Farr, Emilio Gallicchio, Anton Gorenko, Michael M Henry, Frank Hu, Jing Huang, et al. Openmm 8: molecular dynamics simulation with machine learning potentials. The Journal of Physical Chemistry B, 128(1):109–116, 2023.

## References

1. Frey, S., R. Rees, J. Schünemann, S. C. Ng, K. Fünfgeld, T. Huyton, and D. Görlich, 2018. Surface properties determining passage rates of proteins through nuclear pores. Cell 174:202–217.

2. Urry, D. W., D. C. Gowda, T. M. Parker, C. Luan, M. C. Reid, C. M. Harris, A. Pattanaik, and R. D. Harris, 1992.Hydrophobicity scale for proteins based on inverse temperature transitions. Biopolymers 32:1243–1250.

3. Héctor Sánchez-Morán James S. Weltz, D. K. S., and J. L. Kaar, 2021. Understanding Design Rules for Optimizing the Interface between Immobilized Enzymes and Random Copolymer Brushes. ACS Applied Materials and Interfaces

4. Lazaridis, T., and M. Karplus, 1999. Effective energy function for proteins in solution. Proteins: Structure, Function, and Bioinformatics 35:133–152.

5. Jurrus, E., D. Engel, K. Star, K. Monson, J. Brandi, L. E. Felberg, D. H. Brookes, L. Wilson, J. Chen, K. Liles, et al., 2018. Improvements to the APBS biomolecular solvation software suite. Protein science 27:112–128.

6. Mongan, J., C. Simmerling, J. A. McCammon, D. A. Case, and A. Onufriev, 2007. Generalized Born model with a simple, robust molecular volume correction. Journal of chemical theory and computation 3:156–169.

